# ERG activates a stem-like proliferation-differentiation program in prostate epithelial cells with mixed basal-luminal identity

**DOI:** 10.1101/2023.05.15.540839

**Authors:** Weiran Feng, Erik Ladewig, Nazifa Salsabeel, Huiyong Zhao, Young Sun Lee, Anuradha Gopalan, Matthew Lange, Hanzhi Luo, Wenfei Kang, Ning Fan, Eric Rosiek, Elisa de Stanchina, Yu Chen, Brett S. Carver, Christina S. Leslie, Charles L. Sawyers

**Author notes:** These authors contributed equally (Weiran Feng, Erik Ladewig). These authors contributed equally (Nazifa Salsabeel, Huiyong Zhao).

## Abstract

To gain insight into how ERG translocations cause prostate cancer, we performed single cell transcriptional profiling of an autochthonous mouse model at an early stage of disease initiation. Despite broad expression of ERG in all prostate epithelial cells, proliferation was enriched in a small, stem-like population with mixed-luminal basal identity (called intermediate cells). Through a series of lineage tracing and primary prostate tissue transplantation experiments, we find that tumor initiating activity resides in a subpopulation of basal cells that co-express the luminal genes *Tmprss2* and *Nkx3.1* (called Basal^Lum^) but not in the larger population of classical *Krt8*+ luminal cells. Upon ERG activation, Basal^Lum^ cells give rise to the highly proliferative intermediate state, which subsequently transitions to the larger population of Krt8+ luminal cells characteristic of ERG-positive human cancers. Furthermore, this proliferative population is characterized by an ERG-specific chromatin state enriched for NFkB, AP-1, STAT and NFAT binding, with implications for TF cooperativity. The fact that the proliferative potential of ERG is enriched in a small stem-like population implicates the chromatin context of these cells as a critical variable for unmasking its oncogenic activity.

## Introduction

Translocations of the ETS family transcription factor ERG are found in nearly 150,000 newly diagnosed prostate cancers (PCas) each year in the United States^1,2^, underscoring the importance of defining the mechanism by which ERG promotes prostate epithelial transformation. Multiple lines of evidence implicate ERG translocations as an initiating event in prostate cancer. For example, ERG expression is seen in prostatic intraepithelial neoplasia (PIN) and proliferative inflammatory atrophy; expression in cancers is typically uniform; and, in the context of PI3K pathway activation, ERG is sufficient to induce prostate cancer in mice^3–10^.

The mechanism by which ERG initiates prostate cancer remains unclear despite extensive research^5,11–30^. When expressed in primary prostate epithelial cells (in mice or organoids), ERG binds multiple ETS sites throughout the genome but only promotes downstream transcriptional changes in a setting of PI3K activation^5^, which occurs through PTEN loss or PIK3CA family gene mutations in a large fraction of patients with ERG translocations^3,7,31^. Despite these clues, the downstream transcriptional changes responsible for oncogenicity are largely unknown. Bulk RNA-seq analysis of prostate tumors from an autochthonous ERG-driven mouse model, and of ERG+ organoids, has shown that ERG alters androgen receptor transcriptional output and activates a pro-luminal epithelial differentiation program, with consequent loss of basal epithelial cells^5,11,18,32^. These findings raise an intriguing paradox: how does an initiating oncogene that drives terminal epithelial differentiation also promote oncogenesis?

## Results

### ERG stimulates proliferation of epithelial cells with an intermediate lineage state

To gain insight into this question, we re-examined transcriptomic changes in the *Rosa26-ERG^LSL/LSL^; Pten^flox/flox^;Pb-Cre4* (*EPC*) mouse model, now using single cell RNA sequencing (scRNA-seq), and at an early timepoint to capture changes associated with cancer initiation. At age 3 months EPC mice have small foci of invasive cancer which subsequently progress to highly penetrant glandular invasion within 6 months (**Fig. S1A**). Mice with *Pten* deletion only (*Pten^flox/flox^;Pb-Cre4*, hereafter called *PC*) develop intraductal hyperplasia or PIN early and develop invasive cancer only at advanced age (>9 months)^5,33^.

Uniform manifold approximation and projection (UMAP) of all cells from the prostates of *EPC*, *PC* and *WT* mice revealed genotype-specific differences in both epithelial and non-epithelial populations (**Fig. 1A-C, Fig. S1B-F).** For example, *EPC* mice have a pronounced myeloid infiltration not seen in *PC* or *WT* mice (**Fig. S1D-F**)^34^. Focusing initially on the epithelial cells (Epcam+), we noted that the large clusters of secretory luminal cells that account for >90 percent of luminal cells in *WT* mice (called Lum_L1)^35–37^ are absent in *EPC* and *PC* mice. In contrast, a distinct subset of luminal cells (also known as Lum_L2 or LumP), that are localized proximal to the urethra, express stem-like markers (Sca1, Trop2) and have increased regenerative potential^35–37^, are more abundant in *EPC* and *PC* mice (**Fig. 1A-C, Fig. S1G-K**) (hereafter we refer to these Lum_L2-like cells in *EPC* and *PC* mice as Tumor-Lum)^38^. More striking, however, is a cluster of highly proliferative Ki67+, Top2a^high^, Pcna^high^, ERG+ cells with increased S and G2/M cycle scores (cluster 10, **Fig. 1D, Fig. S1L**). Differential gene expression analysis revealed elevated stemness signatures in this proliferative cluster and the adjacent cluster 1, as well as ERG-induced stem, growth factor, and inflammatory signaling, perhaps related to the ERG-associated myeloid infiltrate (**Fig. 1D, Fig. S1M**).

**Fig. 1.**
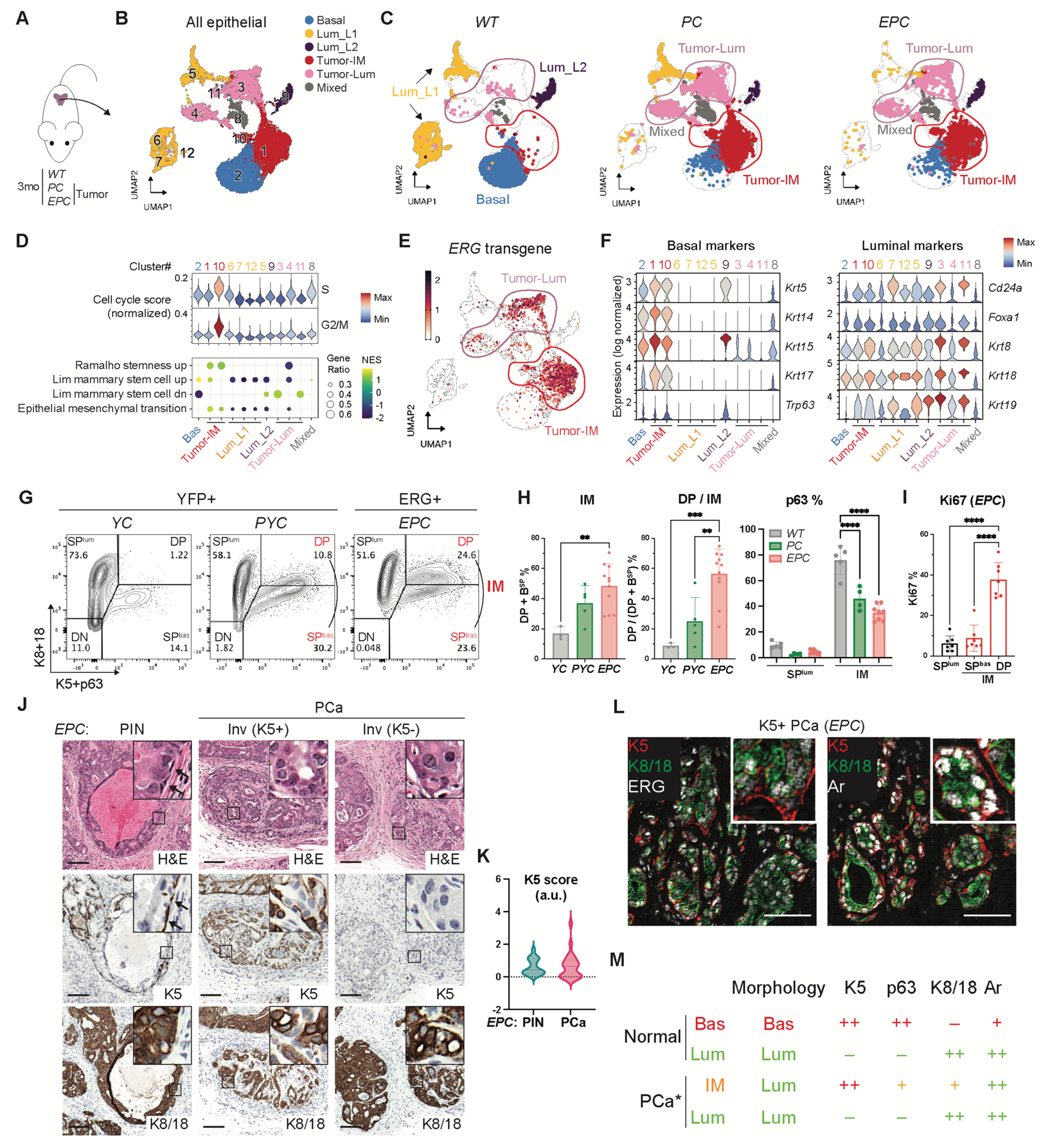
Presence of a highly proliferative intermediate state in ERG-driven prostate cancer. (**A**) Experimental design including *WT* (1 mouse), *PC* (2 mice, *Pten^flox/flox^;Pb-Cre4*) and *EPC* (2 mice, *Rosa26-ERG^LSL/LSL^; Pten^flox/flox^;Pb-Cre4*). (**B-C**) UMAP of epithelial cell clusters with all genotypes together (**B**) or assigned to each genotype (**C**). Luminal and intermediate clusters specific to tumor samples (*PC* and *EPC*) are highlighted in circles and defined as Tumor-Lum and Tumor-IM cells respectively. Cluster IDs are shown. A cluster containing a mix of different cell types is defined as mixed. Bas, basal; Lum, luminal; IM, intermediate. (**D**) Cell cycle scores (top) and gene set enrichment analysis (bottom) across all epithelial clusters. The clusters are numbered according to the UMAP in **B**. Pathways enriched with an FDR-adjusted p-value <0.05 are shown. (**E**) UMAP plotting the *ERG* transgene expression. (**F**) Violin plots comparing canonical basal and luminal marker expression across all epithelial clusters. The clusters are numbered according to the UMAP in **B**. (**G**) Intracellular flow cytometry assessing basal (K5, p63) and luminal (K8, K18) marker expression. Recombined cells labeled by YFP or ERG were analyzed for *YC* (*Rosa26-YFP^LSL/LSL^;Pb-*Cre4)/*PYC* (*Pten^flox/flox^; Rosa26-YFP^LSL/LSL^;Pb-Cre4*) and *EPC* mice, respectively. The combined SP^bas^ and DP fractions in tumors are referred to as IM cells. SP, single positive; DP, double positive. (**H**) Flow cytometry quantification highlighting an expansion of the IM population in tumors and a further luminal shift in IM cells with ERG expression. (**I**) Flow cytometry quantification of Ki67+ cells confirming a high proliferation in the DP fraction of IM cells. (**J**) Haematoxylin–eosin (H&E) prostate histology and immunohistochemistry (IHC) of 3-month *EPC* mice highlighting a luminal morphology of both K5+ and K5-cells from invasive adenocarcinomas, in contrast to the K5+ basal cells that encapsulate the precursor PIN lesions (arrow); inset shows high-power view. PIN, prostatic intraepithelial neoplasia; inv, invasive adenocarcinoma; PCa, prostate cancer. Scale bars, 100 µm. (**K**) Quantification of K5 IHC signal in PIN and invasive adenocarcinomas of *EPC* prostates at 2∼3-month age. (**L**) Immunofluorescence (IF) staining showing expression of ERG (left) and Ar (right) in both K5- and K5+ cells from invasive adenocarcinomas; inset shows high-power view. Scale bars, 50 µm. (**M**) Summary of cell morphology and protein marker expression across different cell types based on flow cytometry and immunostaining in **G-L**. Colors are assigned based on normal basal and luminal cells, which highlight the mixed basal-luminal features of IM cells in PCa. Asterisk, basal cells are absent in PCa, which displays IM cells instead. Data represent mean ± s.d.; n ≥ 3.; **p < 0.01; ***p<0.001; ****p < 0.0001; one-way ANOVA with Tukey posttest.

Due to the luminal phenotype of ERG+ cancers, we anticipated these proliferating cells would be assigned a luminal identity based on their expression signature but were surprised to find instead that these cells were identified as basal, albeit distinct from basal cells in *WT* mice (Tumor-Bas, clusters 1 and 10) (**Fig. S1H**). Importantly, in *EPC* mice these cells express ERG (**Fig. 1E**). Consistent with their basal lineage assignment, cells in Tumor-Bas clusters express canonical basal genes such as *Krt5/14/15/17* and *Trp63*, but also canonical luminal lineage genes such as *Krt8/18/19*, *Cd24a* and *Foxa1* (**Fig. 1F**). Following earlier nomenclature used to refer to rare Krt5/Krt8 “double positive” cells seen in normal prostates^39–46^, we refer to this proliferative population and the adjacent basal-luminal cluster as “intermediate” (IM).

To rule out doublets or other artifactual reasons for co-expression of basal and luminal genes, we sought to confirm the presence of IM cells using flow cytometry. As expected, *WT* mice had two major epithelial populations, single-positive luminal cells (cytokeratin (K)8+/K18+, ∼76%, hereafter called SP^lum^) and single-positive basal cells (K5+/p63+, ∼15%, hereafter called SP^bas^), and a third minor population (∼1%) that was double-positive (DP) (**Fig. 1G**). Consistent with the single cell transcriptomic analysis, *PC* and *EPC* mice had expanded SP^bas^ and DP populations, but with reduced p63 expression relative to *WT* (**Fig. 1G-H, Fig. S2A-B**). In contrast to *WT* mice, where the K5+ population falls almost entirely with the SP quadrant, the K5+ contour plots now overlap with the DP quadrant as a single population. Of note, the contour in *EPC* mice was shifted upward, indicative of higher luminal cytokeratin expression (**Fig. 1G-H**), a finding that we confirmed using a probability-based transcriptomic analysis that assigns quantitative basal or luminal lineage scores to each cell (**Fig. S3**, see methods). Furthermore, roughly 40% of the DP cells in *EPC* mice express high levels of Ki67 (**Fig. I, Fig. S2C**), matching the high proliferation rate seen by scRNA-seq in IM cells. Henceforth, we refer to the entire K5+ population (SP^bas^ + DP) of *PC* and *EPC* mice as IM cells. At a morphologic level, IM cells have a luminal histology and express luminal markers such as K8, K18 and Ar but also express basal genes such as K5 (**Fig. 1J-M**). In addition to K5+ IM cells, a large fraction of ERG+ invasive cells have K5-negative luminal morphology (**Fig. 1J-K**), consistent with the luminal histology of ERG+ human tumors.

### A subset of normal mouse and human prostate basal cells express TMPRSS2

Our single cell analysis of the EPC model revealed that proliferation was restricted to a minor population of epithelial cells with features of both basal and luminal identity (IM). This finding raises the question of the origin of these cells. Does ERG activate a lineage plasticity program in luminal cells that results in mixing of basal and luminal identity, as seen in prostate cancers initiated by loss of the *Trp53* and *Rb1* tumor suppressor genes^47^? Alternatively, does ERG activation occur in a Tmprss2+ cell in the normal prostate with mixed basal-luminal identity that subsequently expands and differentiates into an adenocarcinoma with luminal identity? If so, the fact that ERG activation in prostate cancer is most often a consequence of chromosomal translocation into the *TMPRSS2* locus requires the presence of a TMPRSS2+ cell with mixed basal-luminal identity in the normal prostate.

To explore the latter possibility, we first examined normal prostate scRNA-seq datasets for *TMPRSS2* expression. As expected^48–51^, TMPRSS2 was uniformly expressed in nearly all luminal subpopulations. However, we also noted small numbers of TMPRSS2+ cells residing in basal clusters in both human and mouse normal prostates. Furthermore, these basal cells express other canonical luminal genes such as *NKX3-1*, *NDRG1*, *FKBP5*, *KLK3* and *Pbsn* (**Fig. 2A, Fig. S4A-B**), indicative of a mixed identity. To examine this population more closely, we revisited our previously reported single cell characterization of gene expression changes in the normal mouse prostate during a castration-regeneration cycle^35^. Now, focusing exclusively on the basal cells, we confirmed expression of these same genes in a subpopulation of basal cells as well as their regulation by androgen, just as seen in luminal cells (**Fig. S4C**). To provide additional evidence for luminal gene expression in basal cells, we crossed *Nkx3-1-*CreER^T2^ mice into the *Rosa26-YFP^LSL/LSL^* background. As expected from prior analysis of Nkx3.1 expression (encoded by *Nkx3-1* gene)^52^, the largest population of YFP+ cells were luminal (K8/18+). However, a subset of basal cells (K5+) clearly express YFP (**Fig. 2B**). Importantly, these K5+, YFP+ cells have a flat morphology typical of basal cells yet are adjacent to luminal cells at the basement membrane (**Fig. 2C**). Because these basal cells are single-positive for basal cytokeratin (K5+, K8/18-), and morphologically distinct from the rare, basal/luminal cytokeratin double-positive, triangle-shaped IM cells^39–46^ previously reported in normal prostate (and likely representative of basal cells in transition to luminal cells), we refer to them henceforth as Basal^Lum^ cells (**Fig. 2D**).

**Fig. 2.**
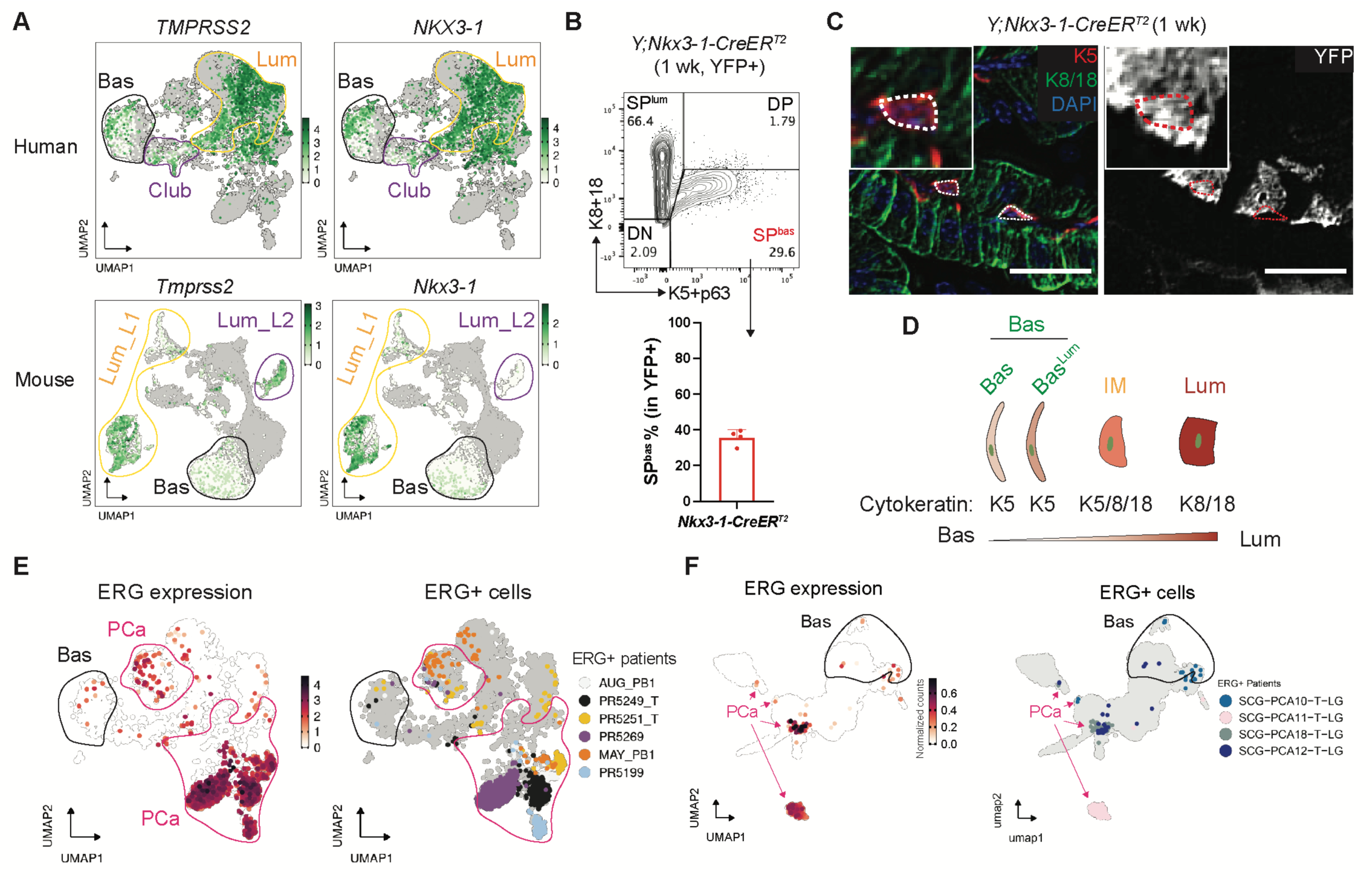
A subset of basal cells express luminal genes in normal prostate. (**A**) UMAP of epithelial cells from normal human and mouse prostates highlighting the expression of canonical luminal genes (*TMPRSS2*, *NKX3-1*, and more in **Fig. S4B**) in a subset of basal cells. Cell types from normal prostates are annotated based on Fig. 1B (mouse) and **Fig. S4A** (human) and highlighted in circles. Cells from normal samples are colored from white to green based on gene expression, with cells from tumor samples in grey in the background. (**B**) (Top) Flow cytometry highlighting the presence of basal cells from the recombined YFP+ population generated by Nkx3-1-CreER^T2^. *Y;Nkx3-1-CreER^T2^* mice were treated with tamoxifen at 2 months and harvested 1 week later. (Bottom) The fraction of basal cells within the YFP+ population are shown (basal proportion may be overrepresented because basal cells survive tissue dissociation better than luminal cells). (**C**) YFP+ cells that co-express K5 from *Y;Nkx3-1-CreER^T2^* mice show a flat basal morphology at the gland periphery; inset shows high-power view. Scale bars, 25 µm. (**D**) Diagram highlighting Basal^Lum^ cells (Bas^Lum^) as a subset of basal cells but distinct from the normal IM cells in cell morphology and cytokeratin expression. (**E-F**) UMAP of epithelial cells from patients highlighting the presence of ERG+ cells in the basal cluster from two independent patient cohorts^53,54^. For each patient cohort, ERG+ cells are colored by ERG expression (left) and patient ID (right). ERG+ PCa samples are shown (see methods). Cell types are annotated based on **Fig. S4A and 5C**, with basal and PCa clusters highlighted in black and pink circles/arrows, respectively.

Having documented that TMPRSS2 is expressed in a rare subset of normal basal cells in mouse and human prostates, we next asked if ERG+ human prostate cancers, which typically display uniform ERG expression in luminal cells, also have evidence of ERG expression in rare basal cells. To address this question, we annotated ERG and TMPRSS2 expression in two independent published scRNA-seq datasets of localized PCa (total PCa n = 29; ERG positive PCa n = 9)^53,54^. As expected, large clusters of ERG+ luminal cancer cells are clearly seen in the ERG+ patients in both datasets, but we also observed rare ERG+ cells in the corresponding basal clusters of these same patients (**Fig. 2E-F, Fig. S5**). Of note, expression of TMPRSS2 in basal cells has also been documented in normal human prostate tissues, primary cultures derived from ERG-positive human tumors, and in mice engineered to express ERG from human *TMPRSS2* BAC construct^55–57^. While larger patient cohorts of scRNA-seq data are needed for a more in depth look at ERG expression in different epithelial cell types, these findings raise the possibility that ERG may initiate prostate cancer in basal cells instead of, or in addition to luminal cells.

### Basal cells are preferred cell of origin for ERG-driven prostate cancer in fresh cell transplantation assays

To explore the cell of origin question, we used a previously reported method to interrogate prostate cancer cell of origin by harvesting primary mouse prostate tissue, sorting into luminal and basal subpopulations, genetically manipulating each subpopulation *ex vivo*, then immediately (same day) reintroducing these cells orthotopically (OT) to score subsequent tumor formation^58^. In our earlier work, luminal cells demonstrated greater tumorigenic potential than basal cells after combined deletion of *Pten* and *Trp53*, confirming results seen previously using luminal or basal-specific Cre drivers^59^. To address the cell of origin using ERG, we isolated luminal and basal cells from *EP* mice, activated the *EP* genotype by infection with Ad-Cre virus and performed OT implantations (**Fig. 3A**). Post-sort analysis confirmed >85% purity of the sorted populations preimplantation and >90% Cre recombination efficiency (**Fig. S6A**). After 5 months, ERG+ cells were detected in both basal and luminal-derived orthografts (**Fig. 3B**) but only the basal-derived orthografts developed invasive adenocarcinomas. Specifically, 6 of 6 mice with basal-derived orthografts developed grossly visible tumors (**Fig. S6B-C**) that contained the highly proliferative ERG+ IM cells and ERG+ SP^lum^ cells (**Fig. 3C**). Histologically however, the basal-derived orthografts displayed luminal morphology, with evidence of nuclear and nucleolar enlargement, and expressed K8 and K18 (as well as K5 in IM cells) with reduced p63. In contrast, luminal-derived orthografts were smaller, less penetrant (4 of 6) and displayed benign histology (**Fig. 3D; Fig. S6B-D**).

**Fig. 3.**
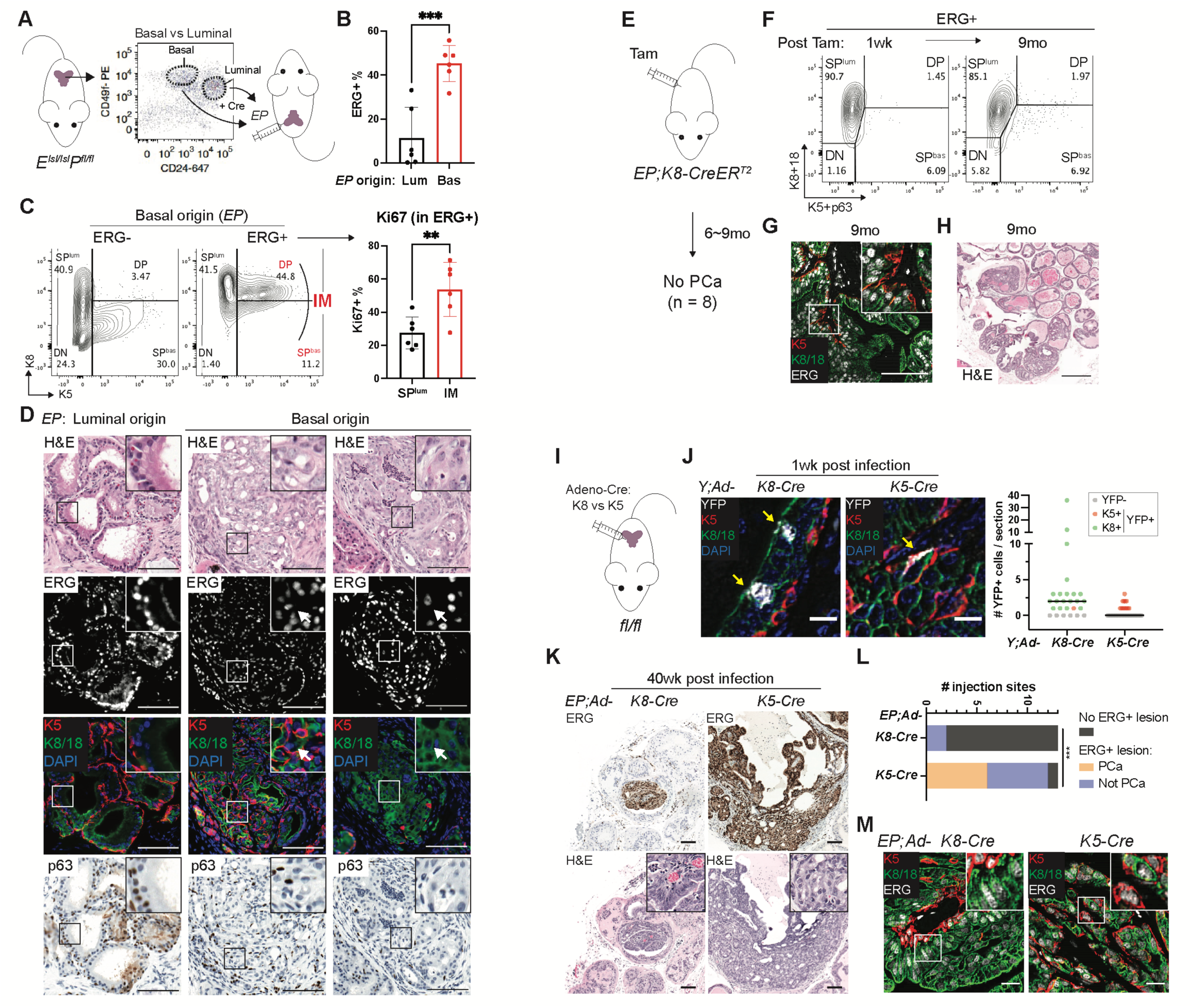
Basal cells are preferred to form ERG-driven prostate cancer with luminal features. (**A**) An orthotopic transplantation approach to compare basal- and luminal-derived *EP* orthografts using freshly isolated and recombined prostate cells. (**B**) Flow cytometry quantification of ERG+ cells in grafts harvested at the 5-month endpoint showing a higher graft burden from the basal-derived orthografts. (**C**) Intracellular flow cytometry highlighting the presence of IM population (left) which were more proliferative than luminal (SP^lum^) cells by Ki67 (right) in ERG+ grafted cells. Mice were harvested at the 5-month endpoint. (**D**) Prostate histological analysis highlighting a histological recapitulation of invasive adenocarcinomas and the presence of luminal cells (K8/18+K5-p63-, arrow) from 5-month basal-originated *EP* grafts; inset shows high-power view. Scale bars, 100 µm. (**E**) A lineage tracing approach to activate *EP* by tamoxifen using a luminal-specific *K8-CreER^T2^* driver. No PCa (defined as invasive adenocarcinoma) was observed after 9 months. (**F**) Flow cytometry highlighting luminal-specific ERG expression from onset (1 week) and maintained through 9 months of tracing in *EP; K8-CreER^T2^* mice. (**G-H**) Prostate histological analysis showing luminal identity (K8/18+K5-) of ERG+ cells (**G**) and lack of PCa development (**H**) in *EP; K8-CreER^T2^* mice up to 9 months of tracing; inset shows high-power view. Scale bars, 100 µm (**G**), 500 µm (**H**). (**I**) An intra-prostatic adenoviral (Ad) injection approach to compare *EP* activation by Ad-K8-vs Ad-K5-Cre. (**J**) IF images (left) and quantification (right) assessing the cell type specificity and infection efficiency of the indicated Ad-Cre at the 1-week time point. Scale bars, 10 µm. (**K-L**) Histology (**K**) and characterization (**L**) of *EP* prostates harvested at 40 weeks post infection; inset, a high-power view highlighting an overall luminal morphology of the invasive cancer cells from *EP;Ad-K5-Cre* mice. PCa is defined as invasive adenocarcinoma. Scale bars, 100 µm. (**M**) IF of *EP* prostates harvested at 40 weeks post infection highlighting the presence of luminal cells (K8/18+K5-) in invasive adenocarcinomas developed from *EP;Ad-K5-Cre* mice; inset shows high-power view. Scale bars, 20 µm. Data represent mean ± s.d.; n ≥ 3.; ***p<0.001; unpaired two-tailed t-test (**B**); Chi-square test (**L**).

### ERG-driven cancers emerge from K5+ but not K8+ positive epithelial cells

Having implicated basal cells as cell of origin in the OT assay, we turned to conventional lineage tracing approaches to validate our findings in an autochthonous model using tamoxifen-inducible *K5-CreER*^T2^ (basal) and *K8-CreER*^T2^ (luminal) mouse strains. Both are well established tools previously used by others to address lineage questions in breast and prostate tissue^60–63^. After confirming the lineage fidelity of these strains in prostate by crossing to a YFP reporter strain (**Fig. S7A**), we generated the relevant *EP* compound mice (*EP*; *K5-CreER^T^*^2^ and *EP*; *K8-CreER*^T2^), administered tamoxifen to cohorts of each strain at age 7-9 weeks, then followed mice for disease onset. Surprisingly, all *EP*; *K5-CreER*^T2^ mice had to be sacrificed within 2-3 weeks of Cre induction due to a fulminant skin disease with extensive desquamation. We observed an essentially identical phenotype in a comparable experiment using an independent basal-specific Cre driver strain (*K14-CreER*^T2^) (**Fig. S7B-G**). This acute skin phenotype was not observed in *Pten^flox/flox^*mice, suggesting that it is a consequence of aberrant expression of ERG in K5/14-expressing cells in skin^64^. Unfortunately, the early death of *EP*; *K5-CreER^T2^* and *EP*; *K14-CreER ^T2^* mice precluded scoring a prostate phenotype.

However, *EP; K8-CreER^T2^* mice remained well following tamoxifen induction and were aged until 9 months. Despite early and sustained luminal-specific ERG induction (at one week, then monthly through the 9 months observation period) (**Fig. 3E-G, Fig. S7H**), we failed to detect any evidence of invasive cancer (n=8 at 6-9 months). The only histologic changes were hyperplasia and PIN in nearly all mice (**Fig. 3H**), a phenotype also seen after Pten loss alone^59,60,65^. We did note rare foci of ERG+ IM cells in a single mouse after 9 months but without evidence of invasive cancer (**Fig. S7I**). Although the *K5-CreER^T2^* experiment was uninformative for scoring a prostate phenotype, the *K8-CreER^T2^* results indicate that K8+ luminal cells are unlikely to serve as cell of origin, consistent with the OT transplantation studies.

To avoid the lethal skin phenotype that complicates use of *K5-CreER^T2^*and *K14-CreER^T2^* mice and directly test whether ERG-driven prostate cancers can initiate from basal cells *in situ*, we delivered Cre locally by intra-prostatic injection of adenoviruses expressing Cre downstream of K5-vs K8-specific regulatory sequences (Ad-K5-Cre, Ad-K8-Cre) (**Fig. 3I**). To evaluate the fidelity and robustness of this method of Cre delivery in the prostate, we used *Rosa26-YFP^LSL/LSL^* reporter mice as a control, as we did previously with the *K5-CreER^T2^* and *K8-CreER^T2^* strains. One week after injection of Ad-K5-Cre, we found all YFP+ cells were K5+ with a flat-shaped basal morphology. In contrast, nearly all YFP+ cells in Ad-K8-Cre injected mice were K8/18+ with a columnar luminal morphology (**Fig. 3J**). Having demonstrated the lineage specificity of the Ad-K5 and Ad-K8-Cre viruses, we performed similar injections in *EP* mice (two injections per mouse into either anterior or dorsal lobes), then followed each cohort for disease onset. In Ad-K5-Cre injected mice, we observed clear expansion of ERG+ cells from 12 out of 13 injection sites, with histologic evidence of invasive adenocarcinoma in half (6 of 12) (**Fig. 3K-L, Fig. S7J**). Despite targeting ERG activation in K5+ basal cells, the ERG+ invasive cells had luminal morphology and expressed K8, K18 and Trop2. ERG+ IM (K5+, Ar+) cells were also observed within the same region (**Fig. 3M, Fig. S7K-L**). In contrast, we did not observe consistent expansion of ERG+ cells in mice injected with Ad-K8-Cre virus (small foci were seen in 2 of 13 mice), and there was no evidence of invasion (**Fig. 3K-M, Fig. S7J-L**). Thus, both the transplantation and *in situ* models converge to the conclusion that basal cells are the preferred cell of origin for ERG-driven prostate cancer, whereas luminal cells support limited, non-invasive cell expansion.

### ERG drives rapid expansion of Nkx3.1+ basal cells

The *Nkx3-1-*CreER^T2^ driver has been used extensively to generate prostate cancer models across a range of genetic contexts (*Pten*^flox/flox^; *Pten^flox/flox^*,*Trp53^flox/flox^*; *Pten^flox/flox^*, *Kras^LSL-G12D^*), often implicating luminal cells as cell of origin for these cancers^65–67^. Having now documented that Nxk3.1 is also expressed in a subpopulation of basal cells^52^, we sought to use this same Cre driver to evaluate the consequence of ERG expression in Nkx3.1+ Basal^Lum^ cells by comparing the relative expansion of these cells after ERG induction versus the more abundant population of Nkx3.1+ luminal cells. Consistent with the YFP data discussed earlier (**Fig. 2**), at one week after tamoxifen induction a small fraction (11%) of ERG+ cells were basal (K5+, K8/18-) whereas ∼85% were luminal. However, a new population of Ki67+, K5+, K8/18+ IM cells emerged one month after ERG induction (**Fig. 4A-C**). Initially, these K5+, ERG+ cells were individual, flat-shaped basal cells adjacent to the basement membrane (one week after induction) but one month later appeared as clusters with a mix of K5+, K8/18-single positive and K5+, K8/18+ DP cells (**Fig. 4D**). After 9 months, all *Nkx3-1-CreER^T2^;EP* mice developed invasive adenocarcinomas. IM populations (K5+, Ar+, morphologically luminal) were evident in 12 of 13 lesions examined, as seen in *EPC* mice (**Fig. 4E-F, Fig. S8A**). Because luminal-restricted ERG expression (*K8-CreER^T2^*) did not initiate invasive cancer (**Fig. 3E-H, Fig. S8B-C**), we conclude that ERG+ cancers preferentially initiate in Nkx3.1*+* Basal^Lum^ cells.

**Fig. 4.**
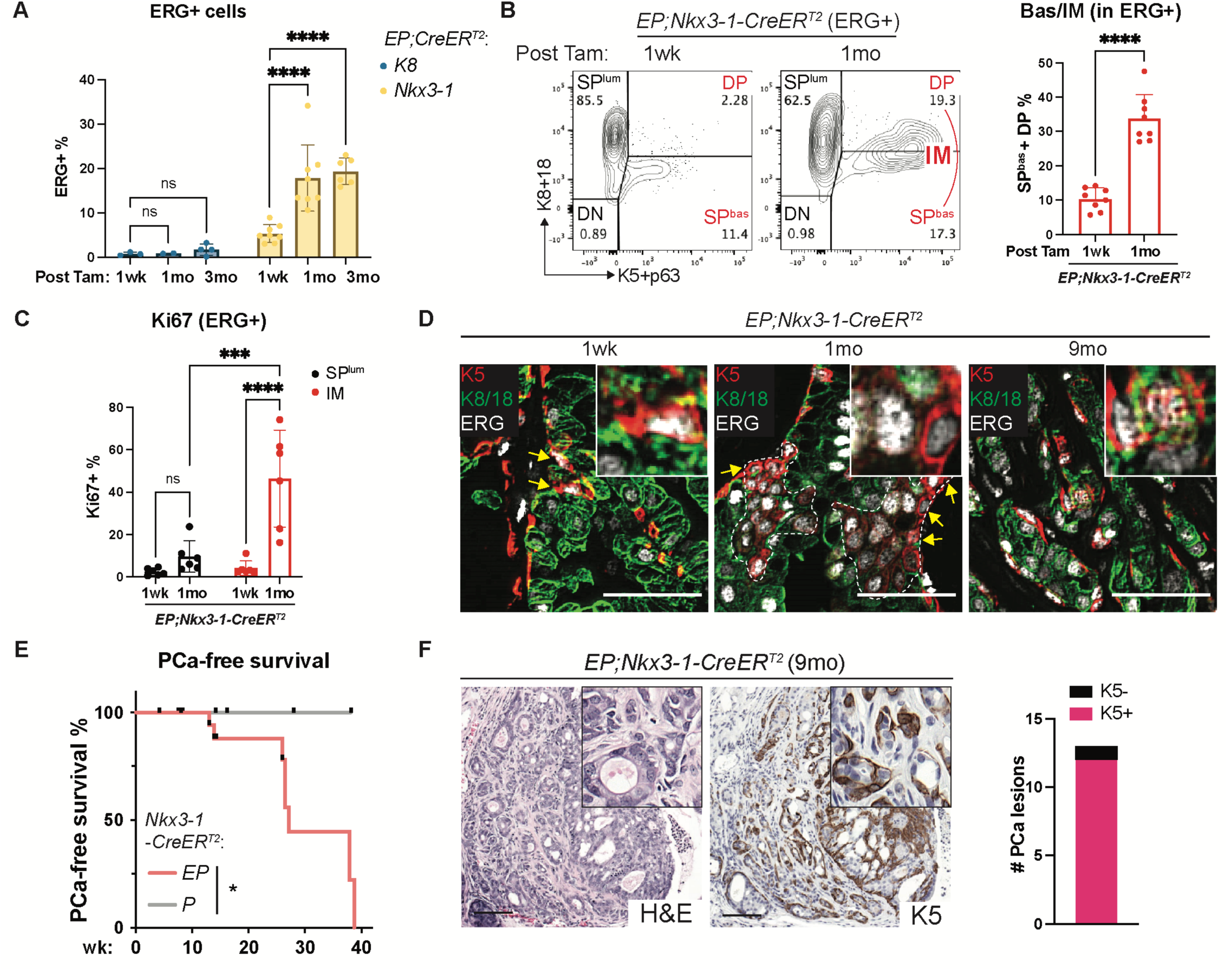
ERG-driven prostate cancer expands from Nkx3.1-expressing Basal^Lum^ cells. (**A**) Flow cytometry quantification of ERG+ cells from *EP* mice activated by the indicated CreER^T2^ drivers along the time course post tamoxifen. (**B**) Flow cytometry from the ERG+ population of the *EP;Nkx3-1-CreER^T2^* mice showing IM expansion by 1 month post tamoxifen. (**C**) Flow cytometry quantification of Ki67+ cells from the ERG+ population of the *EP;Nkx3-1-CreER^T2^* mice harvested at the indicated time points. (**D**) IF highlighting the ERG+ cells that co-express K5 along the time course post tamoxifen. Dashed line encircles clusters of K5+ cells. Arrows highlight the K5-single positive (K8/18-negative) cells at the gland periphery. Inset shows high-power view. Scale bars, 50 µm. (**E**) PCa-free survival analysis. PCa is defined as invasive adenocarcinoma. (**F**) Prostate histology (left) and quantification (right) of K5+ cells in PCas (invasive adenocarcinomas) from *EP;Nkx3-1-CreER^T2^*mice at 9 months post tamoxifen. High-power view (insets) highlights the overall luminal morphology of invasive cells. Scale bars, 100 µm.

### ERG accelerates luminal output from basal cells

Having implicated Basal^Lum^ cells as the source of ERG-driven cancers, we next sought to explain why the histologic phenotype of these cancers is nearly exclusively luminal. First, we used *in vivo* EdU (5-ethynyl-2’-deoxyuridine) labeling of EPC mice with early cancers (age 2 months) to trace the lineage relationships between IM cells and K5-negative, K8+ luminal cells. As expected from their elevated proliferation rate, ERG+ IM cells were preferentially labeled over luminal cells after a brief (2 hour) EdU pulse. However, within 1 week of chase, the label shifted to K8+, K5-negative, ERG+ luminal cells, indicating that ERG+ IM cells give rise to ERG+ luminal cells (**Fig. 5A, Fig. S9A-C**). To provide direct visual evidence for this transition, we identified ∼300 EdU-labeled cell doublets (dividing cells), of which ∼26% showed asymmetric divisions, always with one DP daughter cell paired with a SP^lum^ or SP^bas^ daughter cell, suggestive of a fate transition within one cell division. ∼80% of these asymmetric divisions consisted of DP-SP^lum^ pairs, providing further evidence that DP cells give rise to SP^lum^ cells (**Fig. 5B-C**). Thus, IM cells rapidly transit to a fully luminal state *in vivo*.

**Fig. 5.**
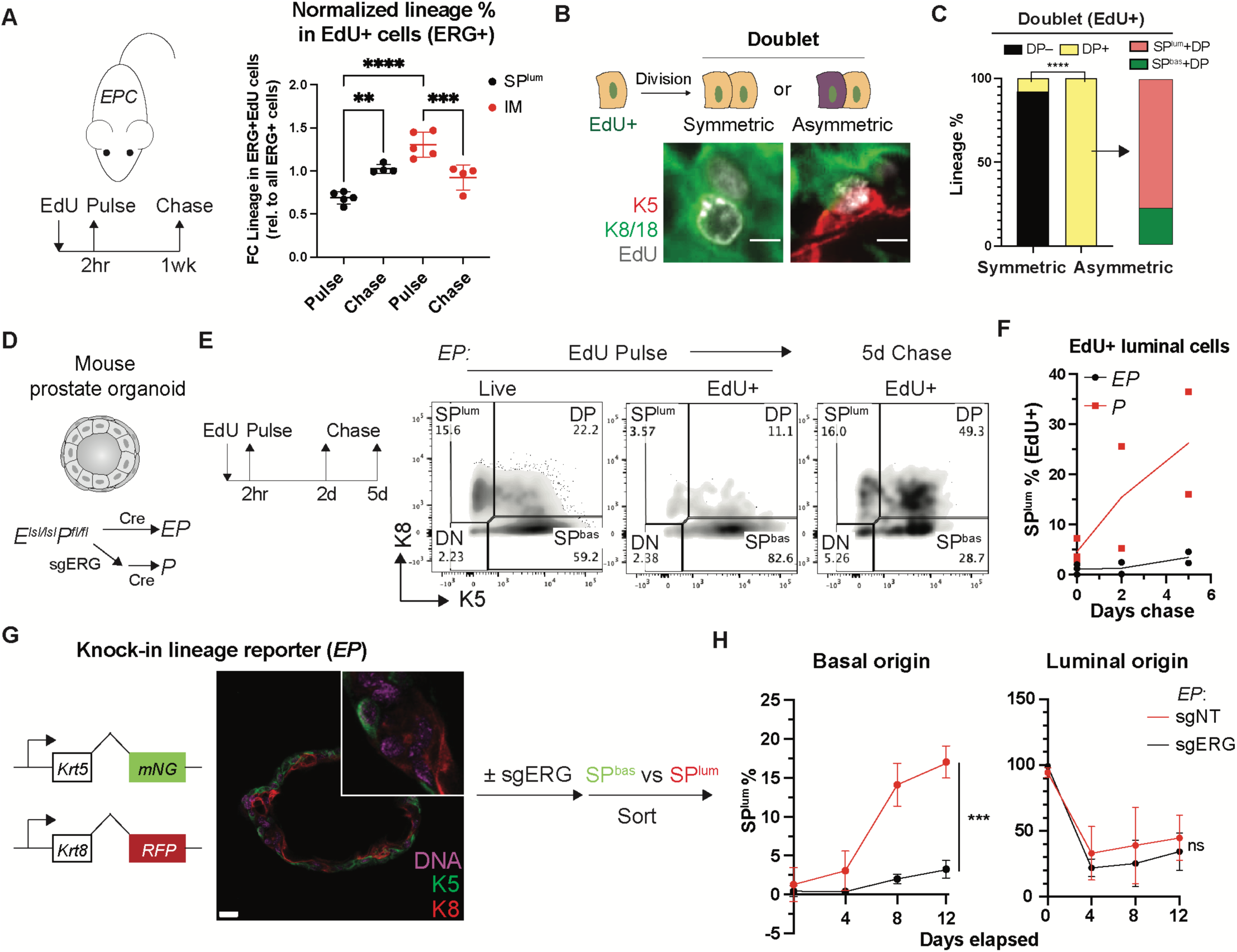
Intermediate and basal cells transit to a luminal fate in the presence of ERG. (**A**) EdU pulse-chase assay on *EPC* mice at 2-month age (schematic on the left). Lineage representation in the EdU+ population of ERG+ cells (normalized to all ERG+ cells) on the right shows an increase of SP^lum^ cells and decrease of IM after the chase. (**B**) Schematic (top) and representative immunofluorescence images (bottom) exemplifying symmetric and asymmetric divisions of EdU-labelled cells. Scale bars, 5 µm. (**C**) The cell type composition of EdU+ doublets highlight that DP cells are present in all asymmetric divisions and preferentially associate with a SP^lum^ daughter cell. (**D**) Schematic showing *in vitro* generation of an isogenic pair of *EP* and *P* organoids. (**E-F**) EdU pulse-chase assay in organoids followed by flow cytometry assessment highlight an ERG-dependent shift of the EdU-labeled cells towards a luminal (SP^lum^) fate. Representative data (**E**) and quantification of the luminal fraction in EdU-labeled populations (**F**) are shown. (**G**) (Left) Schematic showing *EP* organoids with a dual lineage reporter knocked in to form C-terminus fusions with endogenous K5 and K8. (Middle) Live cell imaging highlighting the expected spatial expression pattern of the reporter signals, which enables live sort of basal and luminal cells (schematic on the right). mNG, mNeonGreen; RFP, TagRFP. Scale bar, 20 µm. (**H**) The basal- and luminal-derived *EP* reporter organoids from **G** were analyzed by flow cytometry based on the reporter signals along the time course post sort. An ERG-dependent luminal fate transition from basal cells from luminal cells was observed. Data represent mean ± s.d.; n ≥ 3 (except n = 2 in **F**).; ns, not significant, *p < 0.05; **p < 0.01; ***p<0.001; ****p < 0.0001; one-way ANOVA with Tukey posttest (**A**); Fisher’s exact test (**C**); two-way ANOVA (**H**).

To gain greater clarity into these lineage transitions, we moved from *EPC* mice to a mouse organoid system that allows direct tracing of lineage fate. After establishing primary prostate organoids from *ERG^LSL/LSL^; Pten^flox/flox^* mice, we infected the organoids with Cre virus to delete *Pten* and activate *ERG* expression (**Fig. S9D-E**), then repeated the EdU pulse chase experiment but now *in vitro*. After the 2 hour pulse, the EdU label was primarily in basal cells (>80%) but shifted to DP and SP^lum^ cells during the chase (35% and 27% respectively) and was ERG-dependent (**Fig. 5D-F, Fig. S9F**).

To address whether ERG induction is sufficient to drive a luminal fate transition from purified basal cells, we generated a dual reporter system in *EP* organoids by knocking fluorescent marker genes into the endogenous *Krt5* and *Krt8* loci (**Fig. S10**), thereby providing a platform to isolate live basal or luminal cells for subsequent lineage commitment studies. After confirming the expected localization of the respective K5-targeted (basal) and K8-targeted (luminal) fluorescent signals (**Fig. 5G**), we tracked the lineage fate of sorted basal and luminal cells in organoid culture. ERG-positive basal cells showed a ∼5-fold increase in luminal fate transition compared to ERG-negative cells, confirming that ERG accelerates luminal differentiation from basal cells. In contrast, ERG is not sufficient to sustain luminal fate because sorted luminal cells rapidly decline in relative frequency regardless of ERG state (**Fig. 5H**). Taken together, the EdU and lineage tracing studies support the concept that ERG-driven cancers initiate in rare Basal^Lum^ cells, which expand initially as highly proliferative IM cells then transition to luminal cells.

### ERG establishes a unique chromatin state enriched for STAT and NFAT activity

To gain mechanistic insight into how ERG initiates a program in basal cells that results in proliferation and subsequent luminal lineage specification, we profiled chromatin accessibility changes within the *EPC* model using single cell ATAC sequencing (scATAC-seq). Cell identity of the resulting ATAC clusters was inferred through integration with the scRNA-seq data. Briefly, we aligned scRNA-seq and scATAC-seq datasets using anchor-based integration in order to transfer annotations to scATAC-seq clusters (Methods). To explore the question of how ERG impacts chromatin states, we compared scATAC-seq UMAPs across the three genotypes. Notably, a prominent cluster unique to EPC mice was detected and was transcriptionally defined as IM cells (**Fig. 6A-C, Fig. S11A-G**). Of note, this cluster (hereafter called EPC-IM) is highly enriched for ETS (putatively ERG) binding motifs, implicating ERG as the primary contributor to the shift in chromatin landscape. By contrast, luminal cells of EPC and PC mice (Tumor-Lum) belong to the same scATAC cluster despite the fact that the ETS binding motif is enriched in EPC cells (**Fig. 6A-B, Fig. S11B**). Thus, cell context (IM versus luminal) is critical for ERG to manifest changes in chromatin accessibility. To examine the relationship between cell types, we performed trajectory analysis with Palantir^68^ starting with a cell in the basal cluster based on the lineage tracing data (**Fig. 3**) (Methods). The calculated trajectories suggested some EPC-IM cells precede and may ultimately give rise to Tumor-Lum cells (**Fig. 6D-E**), consistent with the lineage tracing studies described earlier (**Fig. 5A-C**). Together, ERG preferentially changes chromatin landscape in IM cells over luminal cells.

**Fig. 6.**
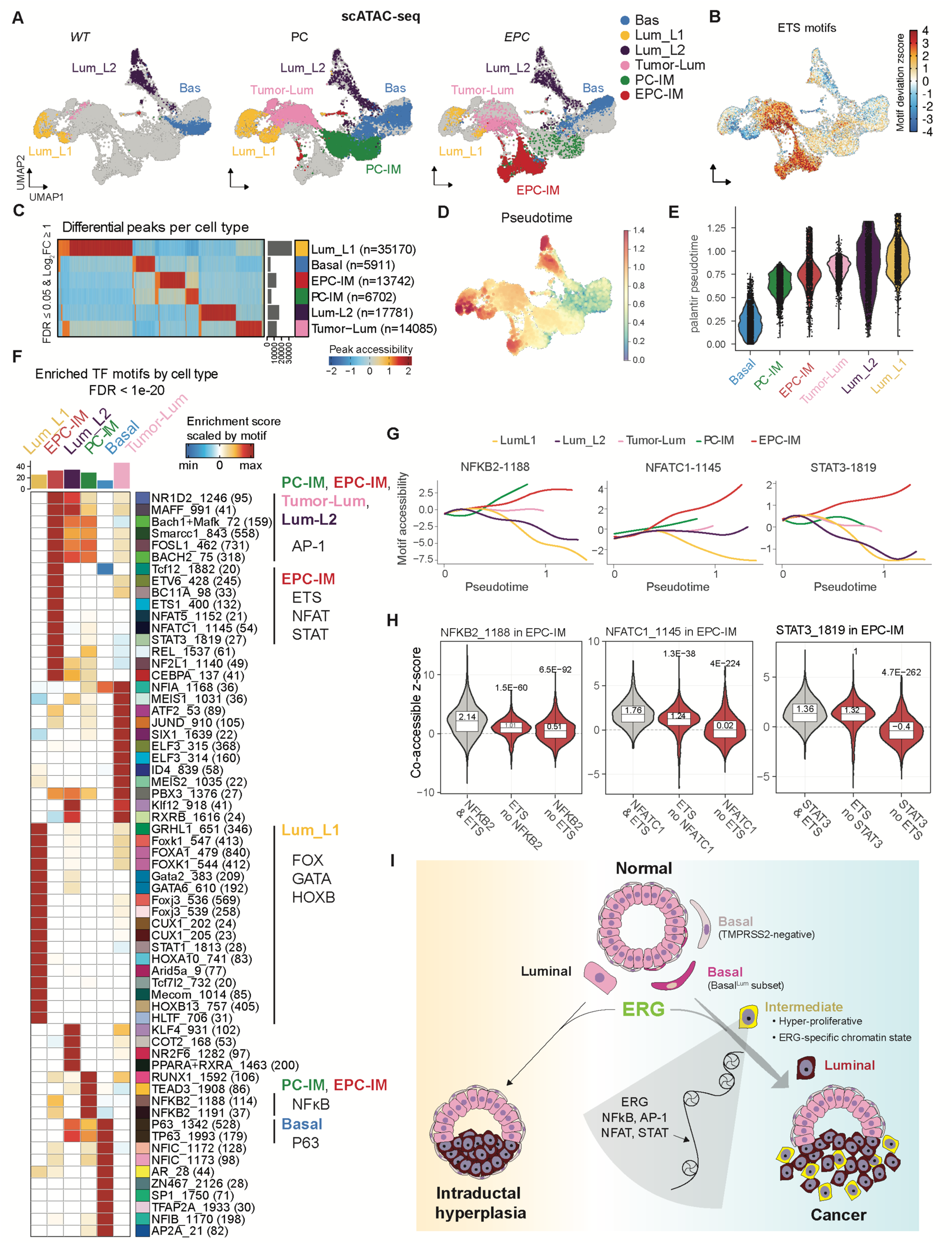
ERG drives a unique chromatin state in intermediate cells. **(A)** Epithelial cell clusters assigned to each genotype (n=16,006 total number of epithelial cells). (B) Single cell chromvar enrichment of ETS family (ERG, ETS, ETV, FLI) motifs. (**C**) Heatmap showing differential peaks between cell types. (**D**) UMAP of cells colored by pseudotime indicating later trajectory branches map to Lum_L1, Lum_L2, and EPC-IM. (**E**) Violin plots showing cells grouped by cell types on the x-axis and arranged by pseudotime on the y-axis. (**F**) Heatmap of transcription factor motifs enriched per cell type. Hypergeometric tests were performed for each motif’s accessibility within each cell type compared to all other cell types. Shown are those motifs having a p-value after false discovery rate adjustment < (10^-20^). Major cell-type specific transcription factor motif families are highlighted. (**G**) Trajectories of 3 selected motifs found with statistically upregulated accessiblity in EPC-IM cells (NFATC1, NFKB2, STAT3). (**H**) Co-accessible chromvar z-scores in EPC-IM cluster of the same motifs shown in (**G**) with or without ETS factors. False Discovery rate adjusted p-values above violin were calculated with 2-sample Wilcoxon rank sum test using chromvar zscores of peaks with co-occurring motifs vs exclusively one motif. (**I**) Model: ERG translocations (in the context of PTEN loss) that occur in a subset of basal cells with luminal transcriptomic features (Basal^Lum^ cells) enter a highly proliferative intermediate state that, in turn drives invasive cancer with luminal differentiation. By contrast, ERG translocations that originate in luminal cells may develop intraductal hyperplasia but fail to progress to invasive cancer. The basal-derived intermediate cells provide a chromatin context to support maximal ERG-driven chromatin changes, featuring motif binding activities of ERG and additional transcription factors (NFκB, AP-1, NFAT, STAT).

The scATAC-seq findings, as well the scRNA-seq and lineage tracing studies described above, collectively point to the EPC-IM cluster as the cell population where ERG activates a tumor initiation program. To gain insight into how this happens, we searched for additional binding motifs enriched within this cluster to identify transcription factors (TF) that might cooperate with ERG. As a quality control, clusters from WT mice with inferred basal and luminal identity showed enrichment of known TF motifs including P63 and FOX/GATA/HOXB13 respectively^48,49,69–74^, confirming the robustness of this approach (**Fig. 6F**). Two patterns emerged in IM clusters: (i) TF motifs enriched in IM clusters from both EPC (EPC-IM) and PC (PC-IM), and (ii) TF motifs enriched in EPC mice alone (**Fig. 6F, Fig. S11H-I**). The first category was noteworthy for enrichment of NF-κB (NFKB2, REL) and AP-1 (FOSL1 and others) family motifs, a finding consistent with previous functional studies linking PTEN loss with elevated NFκB and AP-1 (c-JUN) activity in prostate and other cancers^75–81^. Because PC-IM cells arise earlier than EPC-IM cells in pseudotime, we postulate that PTEN loss leads to activation of NF-κB and AP-1 family TFs in IM cells in both PC and EPC mice.

For the second category of ERG-specific enrichment, we identified STAT and NFAT family TFs in addition to ETS (ETS1, ETV6), all with a high stringency threshold (adjusted p-value < 1x 10^-20^). Based on the correlation of motif accessibility and inferred gene expression, STAT3 and NFATC1 are implicated as the top candidates for each family and, consistently, their motif accessibilities were increasingly enriched across pseudotime in EPC-IM cells, whereas NFκB2 motif enrichment occurs in PC-and EPC-IM cells (**Fig. 6G**). Because STAT and NFAT TFs play critical roles in driving progenitor and inflammatory programs in other tissue contexts including cancer^82–85^, we postulate that STAT3 and NFATC1 may also contribute to the ERG-induced stem and inflammatory programs identified earlier (**Fig. S1M**). It is noteworthy that STAT3 and NFATC1 are both expressed in IM cells regardless of ERG status (i.e. their expression is not induced by ERG) (**Fig. S11I**). However, the accessibility of sites with ERG and additional IM TFs (NFκB2/NFATC1/STAT3) is significantly higher than either TF alone, implicating cooperativity for enabling accessibility (**Fig. 6H**).

## Discussion

ERG translocations are the presumed driver event in nearly half of prostate cancers in Western cohorts (∼150,000 new cases per year), yet we have limited insight into precisely how ERG initiates prostate cancer. This lack of understanding is, in part, due to limited availability of human prostate cancer cell lines, patient-derived xenografts or organoid models that retain ERG expression. Here we have addressed this challenge through single cell interrogation of a GEMM model previously developed by our group that accurately models ERG-driven disease initiation and progression with a histologic phenotype that closely mirrors the human counterpart. This analysis allowed us to identify, unexpectedly, a subfraction of highly proliferative epithelial cells with basal and luminal features (IM cells) that appear within weeks of ERG activation and give rise to the invasive luminal adenocarcinomas that represent the clinical manifestation of the disease. Through a combination of lineage tracing experiments coupled with cell type-specific activation of ERG in basal versus luminal cells (using orthotopic transplantation assays and lineage-specific Cre drivers), we show that ERG-driven prostate cancers initiate in a rare subset of basal cells present in normal mouse and human prostates (which we call Basal^Lum^ cells) that co-express various canonical luminal lineage genes including, importantly, TMPRSS2. Upon ERG activation, Basal^Lum^ cells give rise to the highly proliferative IM cells, which subsequently differentiate into invasive luminal adenocarcinomas (**Fig. 6I**). These findings help resolve and refine preexisting views on the cell of origin of prostate cancer^52,59,60,65,86–91^.

In addition to clarifying the cellular details of how ERG-driven prostate cancers arise, our work identifies the subset of ERG-expressing cells within the tumor (IM cells) that are most relevant for unraveling the molecular details of how ERG drives a program of proliferation and invasion. Toward that end, scATAC-seq analysis revealed a novel, ERG-specific chromatin state exclusively in IM cells and implicates NFkB, AP-1, STAT3 and NFATC1 as TFs that cooperate with ERG in prostate oncogenesis (**Fig. 6I**). NFkB, AP-1 and STAT3 has well established role in oncogenesis across many cancer types, including lung, breast and prostate cancer^83–85,92^. NFATC1 is more closely linked with inflammatory pathways but has been recently implicated in lineage plasticity states in prostate cancer (Zaidi et al., in revision). Additional mechanistic insight into ERG-driven carcinogenesis is likely to arise from deeper interrogation of this IM population, particularly through selective functional perturbations using the *in vivo* models of ERG-dependent oncogenicity described here.

The fact that ERG requires such precise cellular context to initiate cancer presents an interesting contrast with other oncogenic transcription factors such as MYC that act broadly regardless of cell type or tissue. ERG biology in normal tissues provides some precedent. The only tissue that displays absolute ERG-dependence is vascular endothelium, with a phenotype of embryonic lethality in conditional knockout mice due to defects in angiogenesis and blood vessel integrity^93–96^. ERG also plays a context-specific role in hematopoietic stem cells (HSC) by coordinating the balance of self-renewal versus differentiation^97–99^. Interestingly, NFATC1 plays a similar role in HSC maintenance and differentiation^100^, perhaps through cooperativity with ERG in a manner that is now co-opted by the ERG-expressing IM cells in our prostate model. The context specificity of ERG also extends to human cancer, where recurrent oncogenic translocations are limited to prostate cancer (TMPRSS2-ERG), acute myeloid leukemia (FUS/TLS-ERG) and Ewing’s sarcoma (EWS-ERG). Ewing’s sarcoma is of particular interest due to challenges in developing ERG-driven GEMMs despite using a range of Cre drivers^101^, again indicative of exquisite context dependence. Our discovery of a novel chromatin landscape in prostatic IM cells following ERG induction provides a mechanism by which ERG might acquire oncogenic potential by creating a cell type-specific chromatin context in which it can now act. Whether ERG alone can shape this context (as a pioneer-like TF) or in partnership with another TFs remains to be determined.

Our finding of a restrictive cell context for initiating ERG-driven prostate cancer in the mouse raises intriguing questions about the human context, where translocations into the *TMPRSS2* locus can presumably occur in relatively rare Basal^Lum^ cells as well as in highly abundant canonical luminal cells. It is widely accepted that ERG status alone is not prognostic^9,102–106^; however, our mouse data suggest a context where ERG could be prognostic depending on the cell type in which the initial translocation arises. Specifically, translocations arising in Basal^Lum^ cells would be expected to give rise to high grade adenocarcinomas, whereas those arising in canonical luminal cells might be unproductive or generate lower grade carcinomas, as seen in other epithelial malignancies such as lung, breast, and pancreatic cancers with different oncogenic drivers^107–114^. This question is challenging to address using current clinical datasets but should be possible using larger patient cohorts in which the initiating cell can be inferred through single cell analysis.

In closing, our work expands the notion that cancers can initiate in a cell with a different lineage identity from the resultant tumor, a concept first established in chronic myeloid leukemia, where the BCR-ABL translocation initiates in a hematopoietic stem cell but the clinical manifestation of the disease is elevated levels of terminally differentiated myeloid cells (neutrophils)^115,116^. Furthermore, our identification of a highly proliferative, stem-like population with a unique chromatin state may provide an opportunity to reveal novel ERG-specific dependencies that could be exploited therapeutically.

## Acknowledgments

We thank the Sawyers lab for valuable critiques and discussions. We appreciate the feedback from Drs. Michael M. Shen and Cory Abate-Shen, and advice from Cory Abate-Shen lab for setting up the intraprostatic adenoviral injection assay. We are grateful to Rona Lester for help in mouse colony management, the Molecular Cytology Core Facility from MSKCC for help with microscopy, the Single Cell Analytics Innovation Lab Core Facility for help with single cell library preparation, the Integrated Genomics Operation Core Facility for next-generation sequencing, and the Flow Cytometry Core Facility from MSKCC for help with FACS experiments.

## Funding

This work is supported by the following funding sources.

Howard Hughes Medical Institute (HHMI) (CLS)

National Institute of Health grants CA193837 (CLS.), CA092629 (CLS), CA224079 (CLS), CA155169 (CLS), 1K99CA276888-01 (WF)

Department of Defense grant W81XWH-19-1-0323 (WF)

Prostate Cancer Foundation 20YOUN22 (WF)

## Author contributions

Conceptualization: WF, CLS

Methodology: WF, EL, HZ, HL, CSL

Investigation: WF, EL, NS, HZ, YSL, AG, ML, WK, NF, ER

Formal analysis: WF, EL

Resources: YC, BSC

Project administration: ES

Funding acquisition: WF, CLS

Software: EL, CSL

Supervision: CSL, CLS

Writing – original draft: WF, EL, CSL, CLS

Writing – review & editing: WF, EL, CSL, CLS

## Data and materials availability

All data are available in the main text or the supplementary materials.

## Methods

### Mouse procedures

Mouse studies were carried out in compliance with Research Animal Resource Center guidelines (IACUC 06–07–012). *Rosa26-ERG^LSL/LSL^*, *Pten^flox/flox^*, and *Pb-Cre4* mice have been previously described^5,117,118^. To assess Cre activity, *Rosa26-YFP^LSL/LSL^* (*Gt(ROSA)26Sor*^tm3*(CAG-*^ *^EYFP)Hze^/J*) mice^119^ were crossed to the desired strains to introduce the Cre reporter allele.

For tamoxifen-inducible Cre recombination, *K8-CreER^T2^*(*Tg(Krt8-cre/ERT2)17Blpn/J*)^62^, *K5-CreER^T2^* (*Krt5^tm1.1(cre/ERT2)Blh^/J*)^62^, *K14-CreER^T2^* (*Tg(KRT14-cre/ERT)20Efu/J*)^120^ or *Nkx3-1-CreER^T2^* (*Nkx3-1^tm4(cre/ERT2)Mms^/AbshnJ*)^52^ mice were crossed to the desired floxed strains and introduced as hemizygous/heterozygous alleles. To induce CreER^T2^-mediated recombination, tamoxifen (Sigma) was suspended in corn oil and injected intraperitoneally to mice at 7-9 weeks of age. For *K8-* and *Nkx3-1-CreER^T2^*, mice were treated with two doses of 3 mg tamoxifen per 25 g body weight on day 1 and 3, respectively. For *K5-CreER^T2^*, mice were administered one dose of 0.5 mg tamoxifen per 25 g body weight. For EdU pulse-chase experiments, 2-month-old mice were treated with EdU (40 mg/kg; Thermo Fisher) by intraperitoneal injection and harvested at 2 hours (pulse) or 5 days (chase) post administration.

### Histology and Immunostaining

Prostates were collected at autopsy, fixed using 4% paraformaldehyde, dehydrated with 70% ethanol, paraffin embedded, and sectioned on glass slides. Hematoxylin/eosin (H&E) staining was performed following standard protocols by the MSKCC Molecular Cytology Core or IDEXX, Inc. Histological assessments were performed blinded by a MSKCC pathologist.

For immunohistochemistry, paraffin-embedded tissue sections were cut at 5 μm and heated at 58°C for 1 hr. Samples were loaded into Leica Bond RX and pretreated with EDTA-based epitope retrieval ER2 solution (Leica, AR9640) for 20 mins at 95°C. Primary antibodies were applied for 60 mins and detected with Polymer Refine Detection Kit (Leica, DS9800). For detection of P63, an additional incubation with a rabbit anti-mouse antibody linker (Abcam, Cat# ab133469, 0.42 ug/ml) was performed. Similarly, for GFP detection rabbit anti-chicken antibody linker (Jackson ImmunoResearch, Cat# 303-006-003, 2.6 ug/ml) was used. Bound antibodies were detected using Leica Bond Polymer anti-rabbit HRP, followed by Refine Detection Kit Mixed DAB for 10 mins, and Refine Detection Kit Hematoxylin counterstaining for 20 mins. After staining, sample slides were washed in water, dehydrated using ethanol gradient (70%, 90%, 100%), washed three times in HistoClear II (National Diagnostics, HS-202), and mounted in Permount (Fisher Scientific, SP15). Antibody concentrations and catalog numbers are listed in the table below. All FFPE stained tissue was scanned using a MIRAX scanner.

For multiplex immunofluorescence, paraffin-embedded tissue sections were cut at 5 μm and heated at 58°C for 1 hr. Samples were pretreated with EDTA-based epitope retrieval ER2 solution (Leica, AR9640) for 20 mins at 95°C. The staining and detection of multiple antibodies were conducted sequentially. Where applicable, EdU+ cells were detected using Click-iT® Plus EdU Imaging Kit (Thermo Fisher) following the manufacturer’s instructions prior to antibody incubation. For rabbit antibodies, Leica Bond Polymer anti-rabbit HRP secondary antibody was used, for the chicken antibody, a Rabbit anti-chicken (Jackson ImmunoResearch 303-006-003) was used as a linker before the application of the Leica Bond Polymer anti-rabbit HRP. Alexa Fluor tyramide signal amplification reagents (Life Technologies, B40953, B40958) and CF® dye tyramide conjugates (Biotium, 92172, 92174, 96053) were then used for fluorescence detection. After each round of IF staining, epitope retrieval was performed for denaturization of primary and secondary antibodies before another primary antibody was applied. After staining, slides were washed in PBS and incubated in 5 μg/ml 4’,6-diamidino-2-phenylindole (DAPI) (Sigma Aldrich) in PBS (Sigma Aldrich) for 5 min, rinsed in PBS, and mounted in Mowiol 4–88 (Calbiochem). Slides were kept overnight at -20°C before imaging. Antibody concentrations and catalog numbers are listed in the table below. All FFPE stained tissue was scanned using a MIRAX scanner.

| Antibody | Host Species | Concentration/Dilution | Antigen Retrieval | Company | Catalog # | Application |
| --- | --- | --- | --- | --- | --- | --- |
| GFP | Chicken | 2ug/ml | ER2 | Abcam | ab13970 | IHC |
| K5 | Rabbit | 1ug/ml | ER2 | Abcam | ab53121 | IHC |
| K8/18 | Rabbit | 0.12ug/ml | ER2 | Abcam | ab53280 | IHC |
| P63 | mouse | prediluted | ER2 | Ventana | 790-4509 | IHC |
| ERG | rabbit | 1ug/ml | ER2 | Epitomics | 2805-1 | IHC |
| pAkt | Rabbit | 1ug/ml | ER2 | Cell signaling technology | 4060 | IHC |
| Trop2 | Rabbit | 0.5ug/ml | ER2 | Abcam | ab214488 | IHC |
| Nkx3-1 | Rabbit | 0.44ug/ml | ER2 | Proteintech | 13069-1-AP | IHC |
| GFP | Chicken | 2ug/ml | ER2 | Abcam | ab13970 | IF |
| K5 | Rabbit | 0.25ug/ml | ER2 | abcam | ab53121 | IF |
| K8/18 | Rabbit | 0.03ug/ml | ER2 | Abcam | ab53280 | IF |
| Ar | Rabbit | 0.5ug/ml | ER2 | Abcam | ab108341 | IF |
| ERG | Rabbit | 0.5ug/ml | ER2 | Epitomics | 2805-1 | IF |
| pAkt | Rabbit | 0.5ug/ml | ER2 | Cell signaling technology | 4060 | IF |

### Tissue dissociation, organoid culture, and fresh isolation of prostate epithelial cells

Mouse prostates were isolated as described previously^121,122^. Briefly, prostates were harvested and subsequently digested with collagenase type II (Gibco) for 2 hrs at 37 °C, followed by TrypLE (Gibco) digestion at 37 °C until a single-cell suspension was obtained. Digestions were supplemented with Y-27632 (10 μM) to inhibit anoikis and filtered to obtain single cell suspension.

Murine prostate organoid culture was established and maintained under standard conditions as described previously^121,122^. Briefly, dissociated prostatic epithelial cells were embedded in 20-µl drops of Matrigel (Corning) and overlaid with mouse prostate organoid medium. Organoids were passaged by mechanical splitting by repetitively passing through a 200-µl pipette tip or by trypsinization using TryplE. For lineage marker analysis, organoids were trypsinized into single cells and seeded at 20,000-50,000 cells per well of a 12-well plate in the absence of EGF (counted as day 0). For EdU pulse-chase experiments, the organoid culture was incubated with EdU (10 µM, Theremo Fisher) for 2 hrs before harvesting (pulse) or wash-off and re-seeding (chase). To compare cell fate from reporter cells, basal and luminal cells were sorted based on single-positive reporter signals and seeded at equal cell number between sgNT and sgERG in the absence of EGF (counted as day 0). Basal cells were seeded at 20,000-50,000 cells per well of a 12-well plate; less luminal cells were seeded (5,000-10,000 per well) due to a low recovery from the sgERG group.

To isolate basal and luminal cells from donor prostates^123^, freshly dissociated prostate cells were stained for 1 h on ice with CD49f-PE (1:200; BD, 555736) and CD24-Alexa Fluor 647 (1:200; BioLegend, 101818). Sorted basal and luminal cells were transduced with Cre-expressing adenovirus (Ad-CMV-iCre, Vector Biolabs) and directly used for downstream experiments.

To induce Cre recombinase expression, adenoviral transduction was performed as described previously^122^. Briefly, 50,000 dissociated organoid cells or sorted fresh tissue cells were mixed with 1 µl Ad-CMV-iCre (1.0x10^7^ pfu/µl, Vector Biolabs) in 500 µl growth medium. The suspension was spin-infected at 600 g for 1hr before a PBS wash and collecting the infected cells for downstream experiments. The amount of virus used was scaled according to the number of cells available.

### Orthotopic transplantation and intraprostatic adenoviral infection

For orthotopic transplantation of freshly isolated prostate cells, 250,000-450,000 sorted cells were resuspended in 20 μL of 50% Matrigel (Corning) and 50% organoid culture medium before injection into prostate anterior lobes of immunodeficient NSG mice (The Jackson Laboratory) at 2 months of age. Mice were harvested at 5 months post transplantation.

For intraprostatic adenoviral administration, the approach was adapted from previous reports^124–126^ with modifications. Ad-K5-Cre (Ad5-bK5-Cre) (Dr. Anton Berns, Netherlands Cancer Institute) and Ad-K8-Cre (Ad5-mK8-nlsCre)^127^ were obtained at high titer (1.0x10^8^ pfu/µl) from Viral Vector Core at University of Iowa. Adenovirus were prepared in two ways, mixed with 1/9^th^ volume of either 10 mM CaCl_2_, or 16 µg/ml Polybrene (Millipore Sigma). 10 µl adenoviral solution was injected into either dorsal or anterior lobes of floxed mice at 2-3 months of age.

Similar results were obtained which were pooled together in downstream analysis.

### Organoid engineering

Genetic perturbation in organoids was carried out by CRISPR-RNP as previously described^58^. Briefly, Cas9 protein (IDT) was first mixed with sgRNA (IDT) to form the RNP complex before nucleofection. A total of 1.2 µM cRNP was used per individual sgRNA. 500,000-1,000,000 dissociated organoid cells were resuspended with nucleofection buffer, RNP complexes, and electroporation enhancer (IDT, 1:1 molar ratio to RNP) in a total volume of 100 µL. The cell suspension was transferred to a nucleofection cuvette and nucleofected using Lonza Amaxa Nucleofector II (program T-030). Cells were centrifuged and either seeded for culture.

To generate *EP* organoids, the parental floxed organoids (*Rosa26-ERG^LSL/LSL^; Pten^flox/flox^*) were transduced with Cre-expressing adenovirus (Ad-CMV-iCre, Vector Biolabs), sorted based on GFP signal (linked with ERG expression as ires-GFP) and maintained under a non-EGF condition. To generate isogenic *P* organoids, the ERG transgene was preemptively knocked out by CRISPR-RNP from the same parental floxed organoids before Cre transduction. The GFP-negative population sorted out and maintained under a non-EGF condition. The expected ERG and Pten status were confirmed by intracellular flow cytometry at the protein level.

The knock-in lineage reporters were engineered by CRISPaint as previously described^128,129^. An *EP* organoid line with a stable transduction of lentiCas9-blast (Addgene plasmid #52962)^130^ was used as the starting material. To knock in TagRFP at the *Krt8* locus, a targeting vector pCRISPaint-TagRFP (Addgene plasmid # 67167) was electroporated into the organoids together with CRIPSR-RNPs directed to the targeting vector (sgFrame) and *Krt8* (targeting exon 9). The TagRFP+ population was enriched by sorting and prepared for the next steps of engineering. To knock in mNeonGreen at the *Krt5* locus, a targeting vector pCRISPaint-mNeonGreen was electroporated into the organoids together with CRIPSR-RNPs directed to the targeting vector (sgFrame) and *Krt5* (targeting exon 9). The TagRFP-mNeonGreen+ population was first enriched by sorting before a secondary enrichment for the TagRFP+mNeonGreen+ population. The resulting population was used for further study without picking single clones. The knock-in products were first verified by PCR across the break junction (oligos 1 and 2 for Krt8-TagRFP; 3 and 4 for Krt5-mNeonGreen) and Sanger sequencing using the PCR primers. A further verification was performed by intracellular flow cytometry comparing the expression pattern of

K8 vs the TagRFP reporter (not possible with K5 due to the epitope ablation by the targeting event). As a further validation of the Krt5-mNeonGreen engineering, Krt5 depletion by siRNA (siGENOME, Horizon Discovery) led to a reduction of the mNeonGreen reporter signal. The expected spatial expression pattern of the reporters were validated by live cell confocal imaging using Leica SP5 or SP8 confocal microscopes.

sgRNA target sequences:

NT (non-targeting control)^130^: CTTCACGCCTTGGACCGATA

ERG-1^58^: CTGCCGTAGTTCATCCCAA

ERG-2^62^: ATCCTGCTGAGGGACGCGT

Frame^128^: GGGCCAGTACCCAAAAAGCG

Krt8: GATGTCGTGTCCAAGTGAA

Krt5: CTCTTGAAGCTCCTCCGGG

Oligo sequences:

Oligo 1: GTCTCTACCCCTTGTCTCCCT

Oligo 2: CCCTCGACCACCTTGATTCTC

Oligo 3: GCTGTCGTTACAAACAGTGTCTC

Oligo 4: TCCACACCGTTGATGGAGCC

### Intracellular flow cytometry

Intracellular flow cytometry was performed using a Fixation/Permeabilization Solution Kit (BD 554714). Dissociated prostate or organoid cells were fixed and permeabilized with the fixation/permeabilization buffer for 30 min on ice, followed by incubation with primary antibodies for at least 1 hr at room temperature and secondary antibodies (for nonconjugated primary antibodies) for 30 min at room temperature with washes between the steps before analysis by flow cytometry (BD Fortessa or MACSQuant 16). To exclude dead cells, cells were either incubated with live/dead fixable dyes (Thermo Fisher) on ice for 15 min prior to fixation and gated for the negative population, or co-stained with DAPI during antibody incubation and gated by excluding the sub-G1 population. Where applicable, EdU+ cells were detected using Click-iT® Plus EdU Flow Cytometry Assay Kit (Thermo Fisher) following the manufacturer’s instructions prior to antibody incubation. Antibody concentrations and catalog numbers are listed in the table below.

| **Antibody** | **Host Species** | **Dilution** | **Company** | **Catalog #** | **Primary/Secondary** |
| --- | --- | --- | --- | --- | --- |
| K8-AF405 | Rabbit | 1:500 | Abcam | ab210139 | Primary |
| K8-AF647 | Rabbit | 1:500 | Abcam | ab192468 | Primary |
| K18-Biotin | Rabbit | 1:500 | Abcam | ab27553 | Primary |
| K5-APC | Rabbit | 1:500 | Abcam | ab224984 | Primary |
| K5-PE | Rabbit | 1:500 | Abcam | ab224985 | Primary |
| p63-AF647 | Rabbit | 1:500 | Abcam | ab246728 | Primary |
| ERG | Rabbit | 1:1000 | Abcam | ab92513 | Primary |
| ERG-AF488 | Rabbit | 1:500 | Abcam | ab196374 | Primary |
| ERG-AF647 | Rabbit | 1:500 | Abcam | ab196149 | Primary |
| Pten | Rabbit | 1:500 | Abcam | ab170941 | Primary |
| Ki67-PE | Rat | 1:50 | BioLegend | 652404 | Primary |
| CD45-BV605 | Rat | 1:600 | BioLegend | 103139 | Primary |
| CD31-BV605 | Rat | 1:600 | BioLegend | 102427 | Primary |
| TER-119-BV605 | Rat | 1:600 | BioLegend | 116239 | Primary |
| EpCAM-PE/Cy7 | Rat | 1:1000 | BioLegend | 118216 | Primary |
| CD49f-PE | Rat | 1:200 | BD | 555736 | Primary |
| CD24-AF647 | Rat | 1:200 | BioLegend | 101818 | Primary |
| Rabbit IgG | Goat | 1:1000 | Thermo Fisher | A-21245 | Secondary |
| Biotin | Streptavidin-AF405 | 1:1000 | Thermo Fisher | S32351 | Secondary |

### Plasmid construction

To construct pCRISPaint-mNeonGreen, mNeonGreen coding sequence was synthesized and cloned into pCRISPaint-TagRFP (Addgene plasmid # 67167)^128^ between BamHI/NotI sites to replace the TagRFP cassette. The correct cloning is confirmed by Sanger sequencing.

### Sample preparation for single-cell analysis

For scRNA-seq, whole mouse prostates from indicated age were freshly dissociated (described above), and flow sorted for live single cells using DAPI, before processing for scRNA-seq using 10X Genomics Chromium following manufacture’s instructions. For scATAC-seq, dorsal-lateral lobes of mouse prostates from indicated age were freshly dissociated (described above), and flow sorted for live single cells using DAPI. Fresh nuclei were isolated before processing for 10X Genomics Chromium single cell multiome ATAC + gene expression sequencing following manufacture’s instructions. ATAC data were retrieved for downstream analysis.

### Single cell RNA sequencing processing and filtering

scRNA-seq was pre-processed using 10X Cell Ranger software. This included genome alignment and minimal cell filtering for RNA-seq. We used a modified genome reference to align sequencing reads to the Mus Musculus Genome (mm10) plus an additional sequence for the *ERG* transgene. After alignment the 10X Cell ranger software performs UMI and read filtering to create a filtered cell by gene read count matrix. We examined this filtered count matrix to visualize cell and gene coverage distributions and to gather cell / gene statistics. Based on these metrics several plots visualized and used to remove poor quality cells (those with fewer than 500 transcripts) or putative multiplets (defined as cells containing greater than 100K transcripts). Cells detected with 40% or more mitochondrial reads were removed without further consideration.

### scRNA matrix normalization and variance stabilization

To account for technical variation and normalize expression data the R package *Seurat*^131^ was used on the filtered raw count matrix. SCT transform was run to perform normalization, variance stabilization, and feature selection. We regressed out cell cycle scores and mitochondrial percentage in addition to *Seurat* defaults.

### Dimensional reduction and visualization of scRNA

Dimensionality reduction and uniform manifold and approximation projection (UMAP) was applied to the count corrected single cell read matrix to enable a 2-dimensional cell embedding. A nearest neighbor graph of cells was learned and used as input to the leiden algorithm for cell clustering. Ten different cell clustering resolutions were evaluated and scored by silhouette score, a clustering metric. UMAPs were generated and used to hypothesize cell similarities and identify clusters for differential expression testing.

### Differential genes and pathways

Differential expression testing was performed using the R package *presto* (https://github.com/immunogenomics/presto). Genes were ranked using the average log fold change and a false discovery rate adjusted p-value. A GSEA analysis with fsGSEA^132^ utilized a list of ranked genes per comparison and the collection of molecular signatures database (MSigDB).

### Epithelial cell identification

We investigated all single cells for epithelial cell populations by summarizing clusters of Epcam+ cells. Cell clusters containing 50% or more of Epcam+ cells were flagged as epithelial related. Next individual cells were scored using an epithelial cell signature set of differentially expressed genes defined in wildtype mouse by Karthaus et al.^35^. Those cells found in flagged epithelial clusters and defined as basal, luminal 1 (L1), luminal 2 (L2), or luminal 3 (L3) were retained as confident epithelial cells and used in downstream analysis and clustering.

### Luminal and basal cell probabilities

As an orthogonal approach to epithelial cell identification, we scored individual cells with a set of luminal and a set of basal cell related genes found in literature. These marker genes scores were then used to cluster cells using a multivariate gaussian mixture model. The mixture model identified three distributions that were also easily identifiable in a scatter plot of basal high, luminal high, or high in markers from both basal and luminal genes (termed intermediate, IM). Those cells found to be up in both markers sets were mostly located within a tumor-only cluster and previously identified as epithelial basal by another gene set from Karthaus et al. (Epithelial cell identification section)^35^. Cell probabilities for the three types were calculated from the gaussian mixtures and subsequently used to assign cells to their most probable type, either luminal, basal, or intermediate.

### Human scRNA data

Human primary prostate cancer single cell RNA sequencing data from the publication^53,54^ are accessible from GEO under accession numbers GSE176031 and GSE181294, respectively. Data were reproduced from these studies using the same cell type classification with the following exceptions. For the Song et al. study ^54^, the two subtypes of PCa cells which were originally annotated as ERG-positive and ERG-negative tumor cells are now together named as PCa in this study because ERG+ cells also exist in the “ERG-negative” clusters. The ERG-positive and mixed (denoting those carrying both ERG-positive and ERG-negative tumor cells by scRNA-seq) patients defined by the original study are now grouped together as ERG+ patients in this study considering the typically uniform ERG expression pattern in PCa by histology^4,6,9^. Similarly for the Hirz et al. study ^53^, ERG+ patients were defined by those with >50% PCa cells to be ERG+.

### Single cell Assay for Transposase-Accessible Chromatin with high-throughput sequencing data (scATAC-seq) filtering features matrix

Single cell Assay for Transposase-Accessible Chromatin with high-throughput sequencing data (scATAC-seq) was evaluated from the ARC filtered count matrix. Cell filtering, feature selection, and analyses were performed using *ArchR* software^133^. We generated output plots to visualize distributions of cell counts within promoters, gene bodies, and transcription start sites (TSS). Quality metrics were used to remove poor quality cells including low TSS enrichment (<4) or low number of unique nuclear fragment (<1000) or putative multiplets (defined as cells containing greater than 1M reads).

### Dimensional reduction

Dimensionality reduction was calculated via *ArchR* with Iterative Latent Semantic Indexing using 500bp genomic tiles. A nearest neighbor graph of the LSI reduced data was performed by *Seurat’s* shared nearest neighbor implementation. We excluded the first LSI component that was correlated > 75% to read depth. Clusters were then identified with the leiden algorithm.

### Peak and motif identification

Peaks with scATAC-seq data were called using *MACS2*^134^ through the *ArchR* interface. Differentially accessible regions were defined as those having significant (FDR < 0.05) difference with log_2_ fold change ≥ 0.5. For motif detection we utilized 2 databases, 1) CisBP and 2) the Non-redundant TF motif matches genome-wide^135^. Motif matching within peaks was performed using *MotifMatcher* with defaults. The gene identity of each position weight matrix match was recorded so that we retained only those motifs with corresponding normalized average gene expression per cell type > 1. To identify motifs statistically significant between comparisons we calculated motif *ChromVar*^136^ scores per cell and performed Spearman correlation of those scores to the motif’s gene score values. Those motifs having positive correlation and significant ChromVar scores per cell type were retained.

### Motif co-accessibility

The ETS family of factors includes several highly similar transcription factor motif position weight matrices. To simplify the identification of ETS family factors within scATAC-seq we visualized the distribution of similar ETS family motif deviation z-scores before reducing the following motifs from oncogenic ETS family TFs in prostate cancer^31^: ERG, ETS, ETV, FLI, into one identifier, hereafter referred to as ETS motifs. To evaluate co-accessibility of ETS motifs with other significantly accessible motifs within cell types, called candidate motifs (**Methods**: Peak and motif identification) we identified peaks containing each candidate motif and the ETS motifs. We then calculated the per cell deviation and deviation z-score with Chromvar. Similarly, we calculated chromvar z-scores for each candidate motif whose peaks did not overlap ETS motifs and then the complement, i.e. ETS motifs peaks that do not harbor the candidate motif. For each candidate motif a 2-sample Wilcoxon rank sum test was used to establish whether co-occurance of the candidate and ETS motifs were significantly different (one-sided alternative “greater”) from: ETS motifs that do not co-occur with the candidate motif, or 2. candidate motifs that do not oc-occur with ETS motfs. For NFKB2, NFAC1, and STAT3 the co-occurrence with ETS were significantly greater than without ETS within EPC-IM cells (**Fig. 6H**).

## Supplemental figures

**Figure S1.**
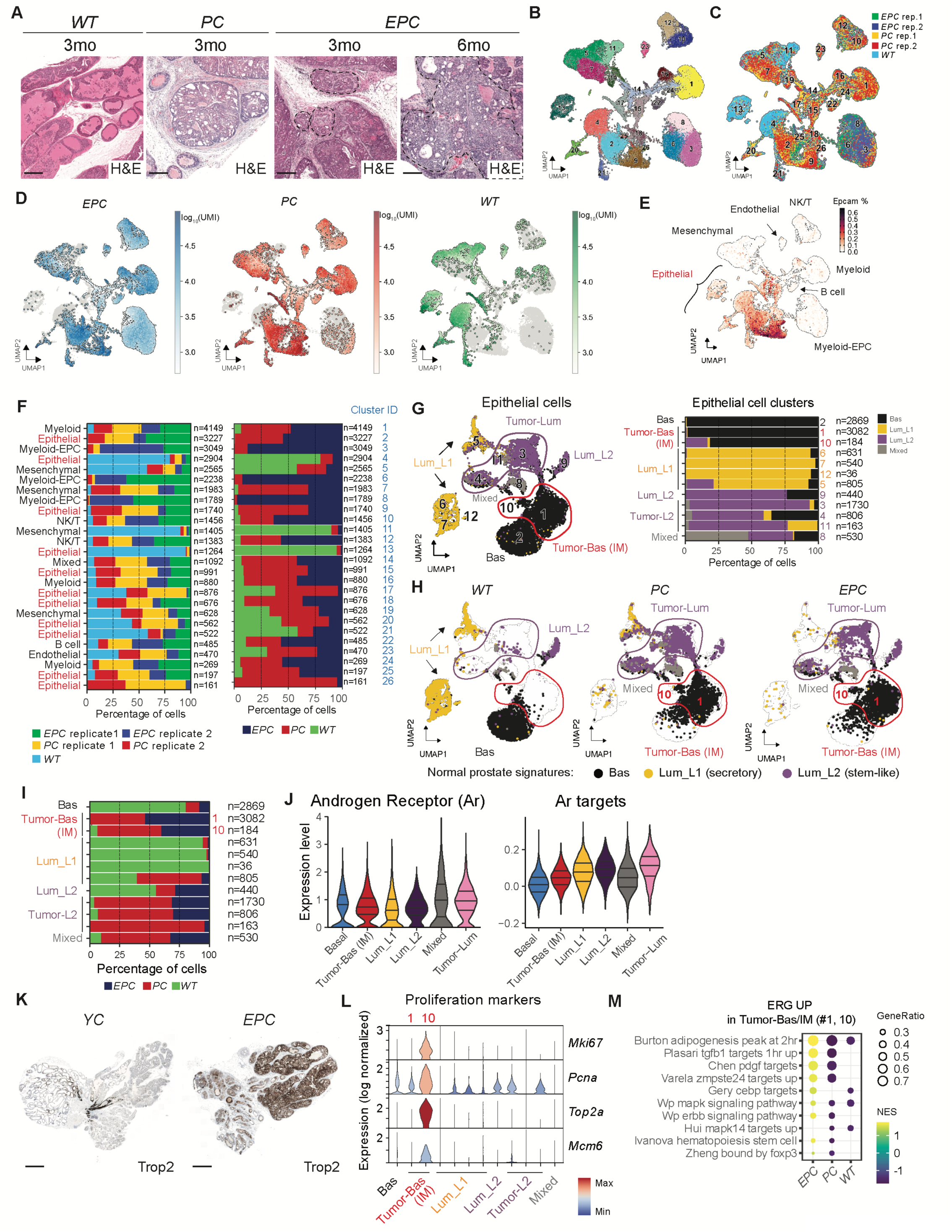
Time point characterization and annotation of scRNA-seq clusters of genetically engineered mouse models (GEMMs). (**A**) H&E prostate histology. Dashed lines encircle invasive adenocarcinoma highlighting an ERG-dependent focal invasion from 3-month *EPC* mice that later became pervasive at 6 months age. Scale bars, 250 µm. (**B**) UMAP of all cells (N = 36,961 cells) from the prostates of 5 mice (2 *EPC*, 2 *PC*, 1 *WT*) annotated based on leiden clustering. (**C**) UMAP of all cells colored by individual replicates per genotype. (**D**) UMAP of all cells showing cells assigned to each genotype and colored by log10(UMI). **(E)** UMAP of all cells previously defined normal mouse prostate signatures. Epithelial cell clusters defined based on *Epcam* expression are highlighted. Myeloid cells specific to the *EPC* samples were defined as Myeloid-EPC. (**F**) Barplot of all cells colored by individual replicates (left) or each genotype with replicates aggregated (right). Numbers of cells in each cluster were included (n=) and cluster number corresponding to **B** is shown. (**G**) UMAP (left) and barplot (right) of all epithelial cells colored by cell types based on normal prostate signatures^35^. Numbers assigned to each individual cluster are shown. Colored circles highlight L2 cells (purple) and basal-like cells (red) specific to *PC* and *EPC* prostates (termed Tumor-Lum and Tumor-Bas respectively). Tumor-Bas cells were later defined as Tumor-IM cells as shown in **Fig. 1B**. A cluster containing a mix of different cell types is defined as mixed. Bas, basal; Lum, luminal. (**H**) UMAP showing epithelial clusters assigned to each genotype. Clusters are annotated as in **G**. The IDs of the two Tumor-Bas/IM clusters are shown. (**I**) Barplot of epithelial clusters showing the cell type composition of each cluster. Numbers of cells in each cluster were included. The clusters are numbered according to the UMAP in **G**. (**J**) Violin plots showing Androgen receptor (Ar) expression (left) and Ar signature scores (right) across epithelial cell types. (**K**) Sagittal view of whole prostates with Trop2 IHC on mice at 3 months age. In a normal prostate (*YC*), Trop2 selectively stains the stem-like L2 luminal cells at the proximal regions and distal invagination tips over the secretory L1 luminal cells, as expected^35,36,137,138^. By contrast, *EPC* tumor cells displayed a pan-Trop2 staining regardless of their tissue localization, further supporting a pervasive L2 transition as revealed from scRNA-seq analysis in **G-H**. Scale bars, 1 mm. (**L**) Violin plots comparing proliferative marker expression across all epithelial clusters. The two Tumor-Bas (IM) clusters are numbered according to the UMAP in **G**. (**M**) Gene set enrichment analysis showing EPC-specific pathways (ERG UP) within the Tumor-Bas/IM population.

**Figure S2.**
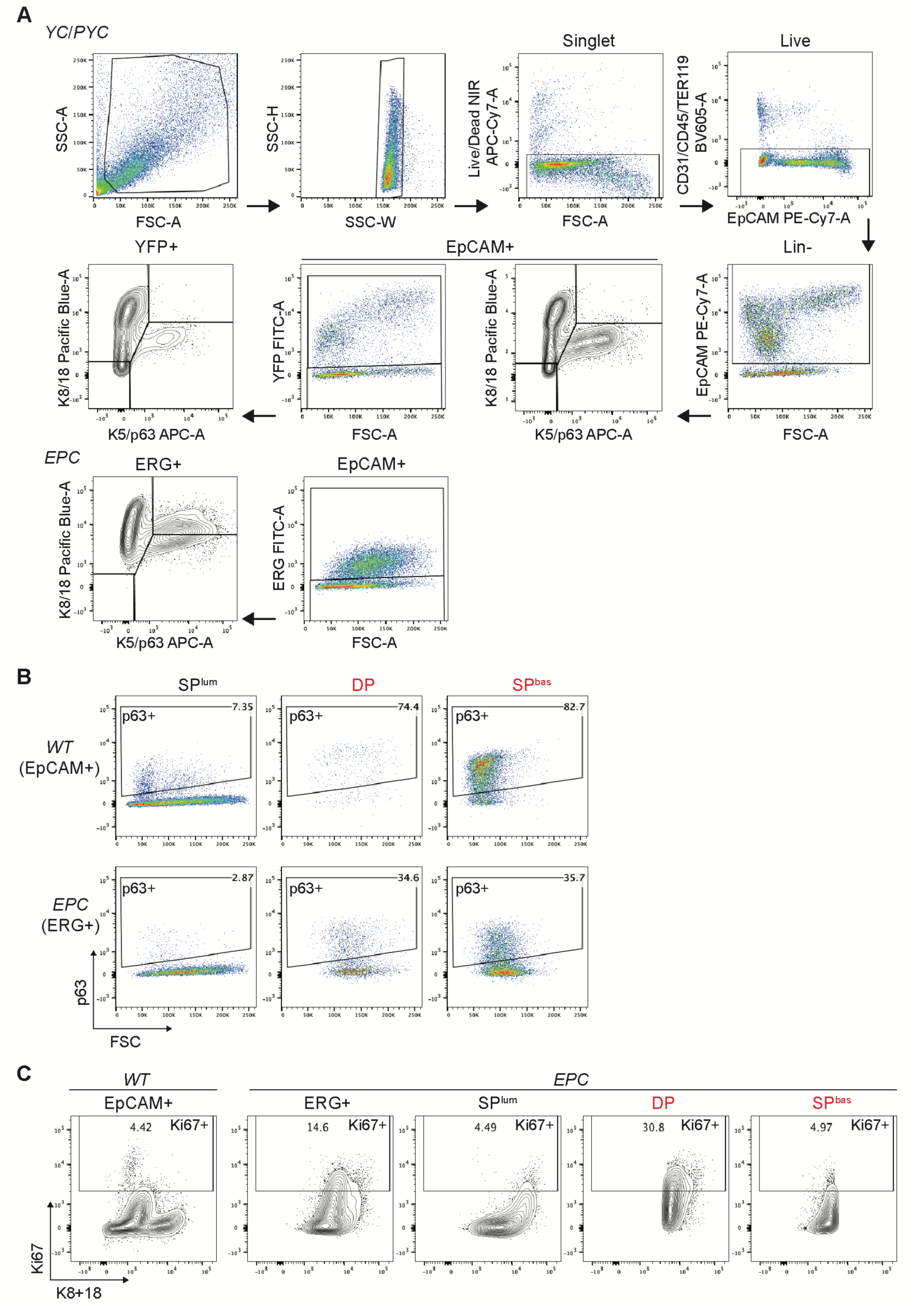
Flow cytometry panel design. (**A**) Single cell suspension of dissociated prostate cells were gated for debris elimination, singlets (SSC-H vs -W), live cells (Live/Dead NIR exclusion), endothelial/immune exclusion (Lin-, CD31-,CD45-,Ter119-) and epithelial cells (EpCAM+). The epithelial population were further gated on the Cre-recombined population (YFP+ for *YC*/*PYC*, ERG+ for *EPC*) before analyzing basal (K5/p63 single positive, SP^bas^), luminal (K8/18 single positive, SP^lum^) and double-positive (DP) populations. DP population is gated as the outlier population based on the *WT* contour plot, assuming that DP cells rarely exist in normal adult prostates. The continuous population of SP^bas^ and DP in tumor samples (*PYC*/*EPC*) are combined and named as IM. (**B**) To assess p63 expression, epithelial or ERG/YFP+ cells were first gated as in **A**, before applying the p63+ gate to populations of interest. (**C**) To assess Ki67 expression, epithelial or ERG/YFP+ cells were first gated as in **A**. Ki67-positive cells were gated as the outlier population based on the *WT* contour plot, assuming that normal adult prostates are rarely proliferative, before applying to other populations of interest.

**Figure S3.**
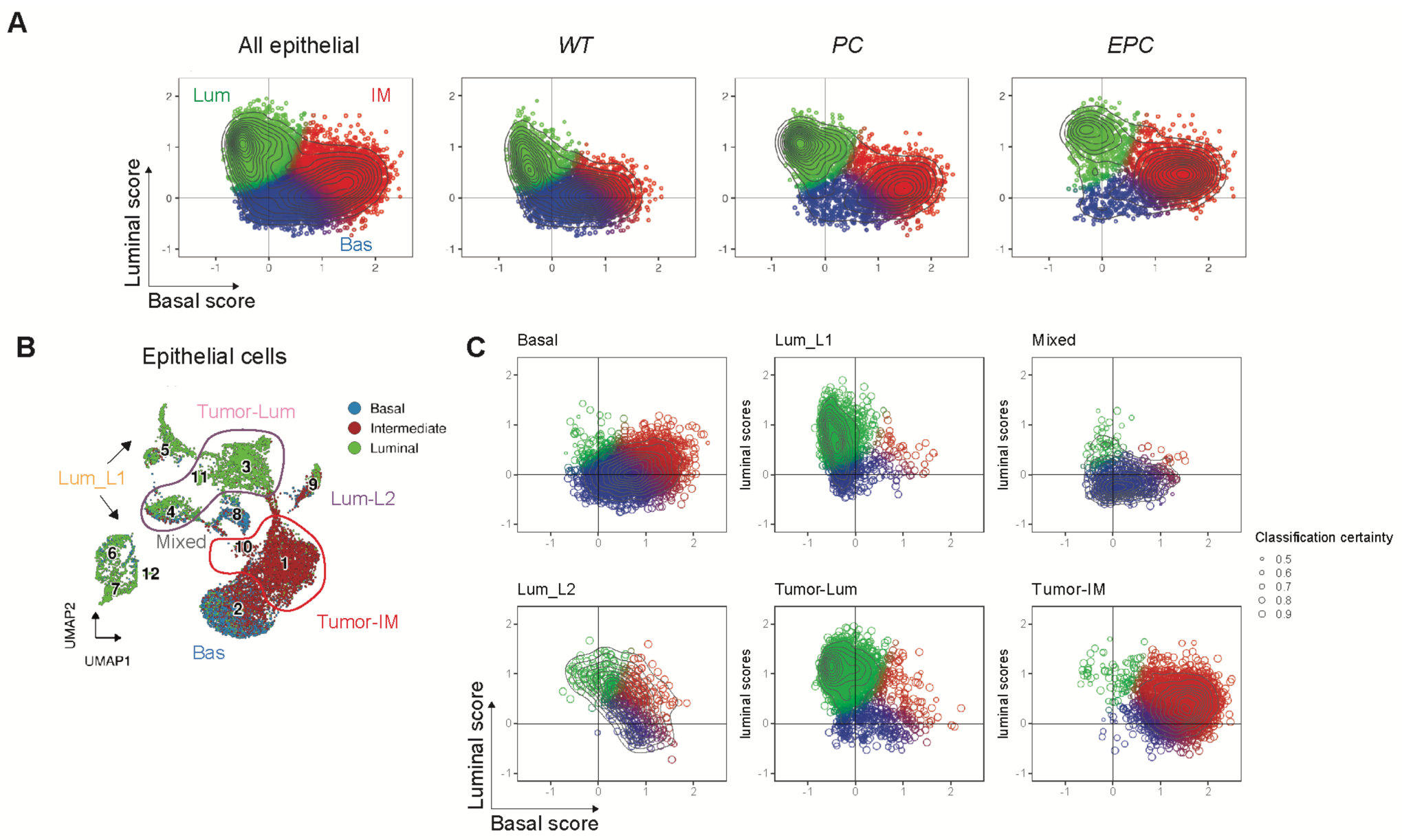
Tumor-IM cells show intermediate features. (**A**) Assignment of basal and luminal scores to epithelial cells from all samples aggregated and each individual genotype (see methods). Unsupervised analysis revealed three distinct populations from all epithelial cells which were named as basal, luminal, and intermediate according to the marker scores. A pronounced double-positive IM population in *PC* and *EPC* mice are shown. (**B**) UMAP of all epithelial cells highlighting the IM identity of the Tumor-IM clusters. Clusters are colored by basal, luminal, and intermediate cell types defined in **A**. (**C**) Assignment of basal and luminal scores in each individual cell type.

**Figure S4.**
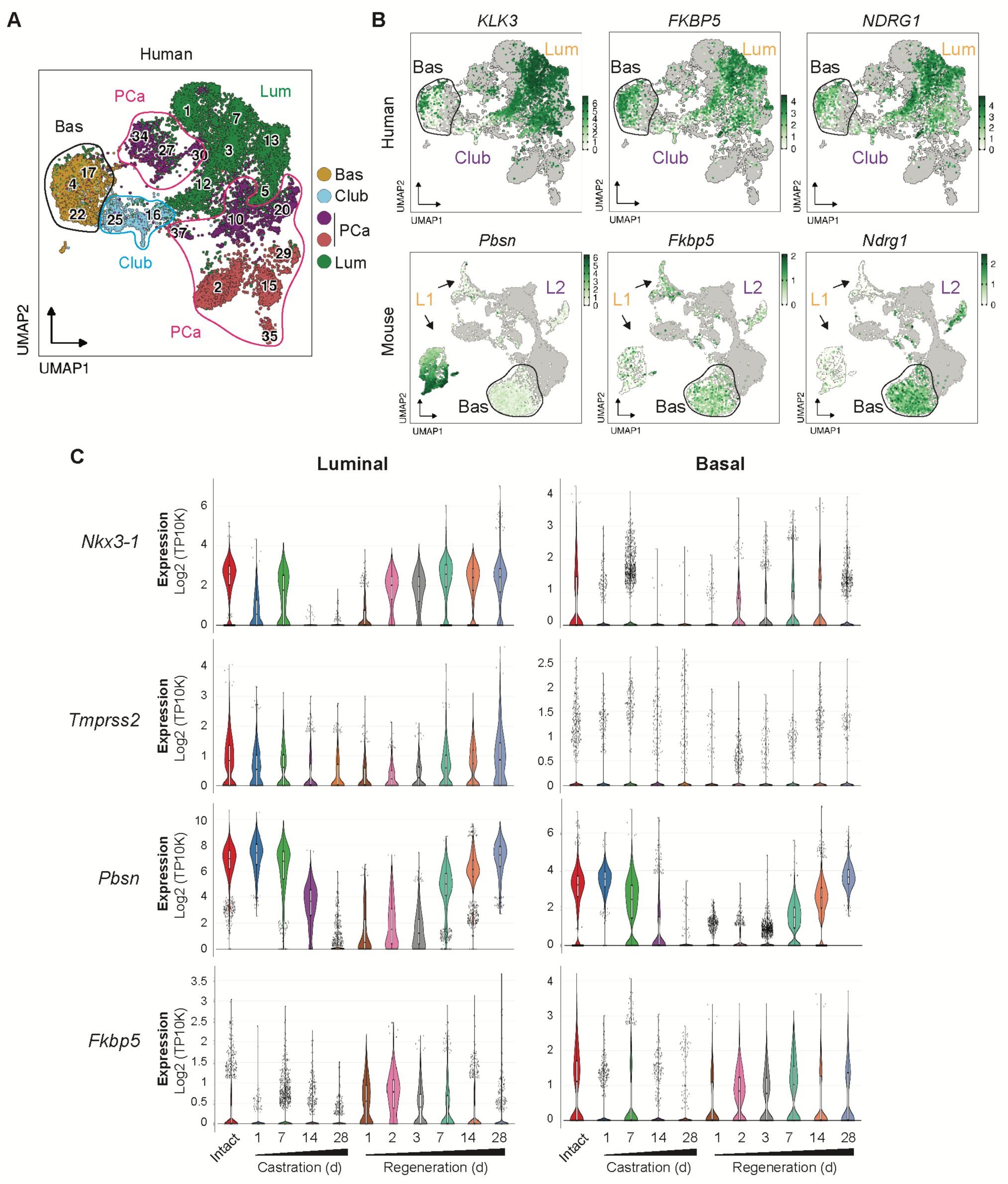
A subset of basal cells express androgen-regulated luminal genes. (**A**) UMAP of all epithelial cells from human prostates. Data were reproduced from a previous study^54^ with the same cell type classification, except that the two subtypes of the previously defined prostate cancer cells are together named as PCa here (see methods). The basal and PCa clusters are highlighted in black and pink circles, respectively. (**B**) UMAP of epithelial cells from normal human and mouse prostates highlighting the expression of canonical luminal genes in a subset of basal cells. An expanded list of genes beyond **Fig. 2A** are shown. Normal basal clusters are highlighted in circles. Cells from normal samples are colored from white to green based on gene expression, with cells from tumor samples in grey in the background. The complete UMAPs and cell type annotations are shown in **A** (human) and **Fig. 1B** (mouse). (**C**) Changes in expression of luminal genes expressed in a subset of basal cells during a castration-regeneration cycle, indicative of androgen dependent expression, as seen in canonical luminal cells. Data were mined from a previous publication^35^.

**Figure S5.**
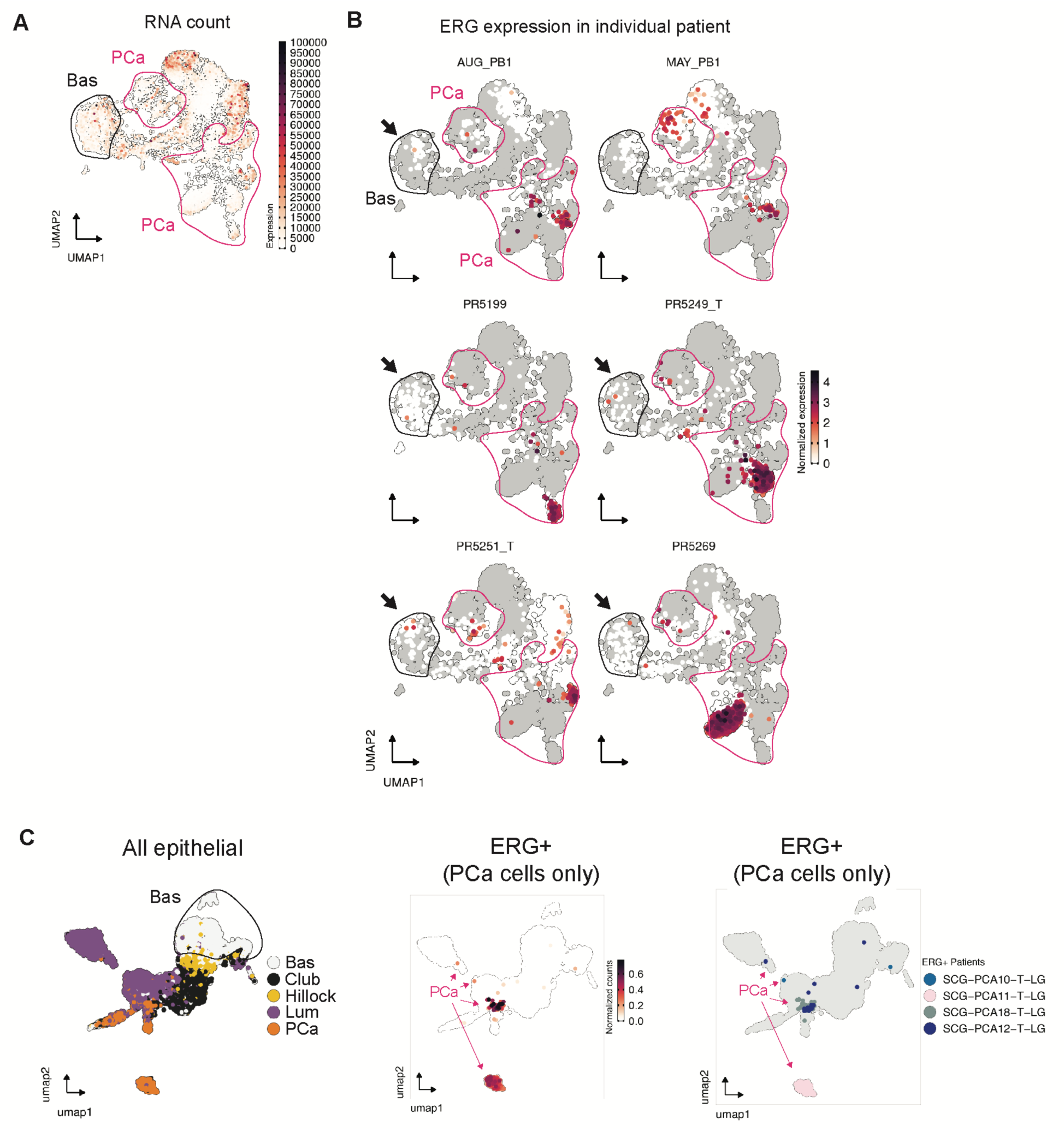
Evidence of ERG+ basal cells in human prostate cancer. (**A**) UMAP of epithelial cells from patients^54^ showing RNA count to assess doublet potential. The low to intermediate level of RNA count in basal cluster suggest the ERG+ basal cells in **Fig. 2E** are unlikely an artificial outcome of doublet formation. (**B**) UMAP of epithelial cells showing ERG expression in individual patients. ERG+ PCa samples are shown (see methods). Cells from each patient are colored from white to red based on gene expression, with cells from the rest of patients in grey in the background. Arrows highlight the presence of ERG+ cells in the basal cluster. Cell types in this figure are annotated based on **Fig. S4A**, with basal and PCa clusters highlighted in black and pink circles, respectively. (**C**) UMAP of epithelial cells from a different patient cohort^53^ showing all epithelial cells (left), ERG+ cancer cells colored by ERG expression (middle) and patient ID (right).

**Figure S6.**
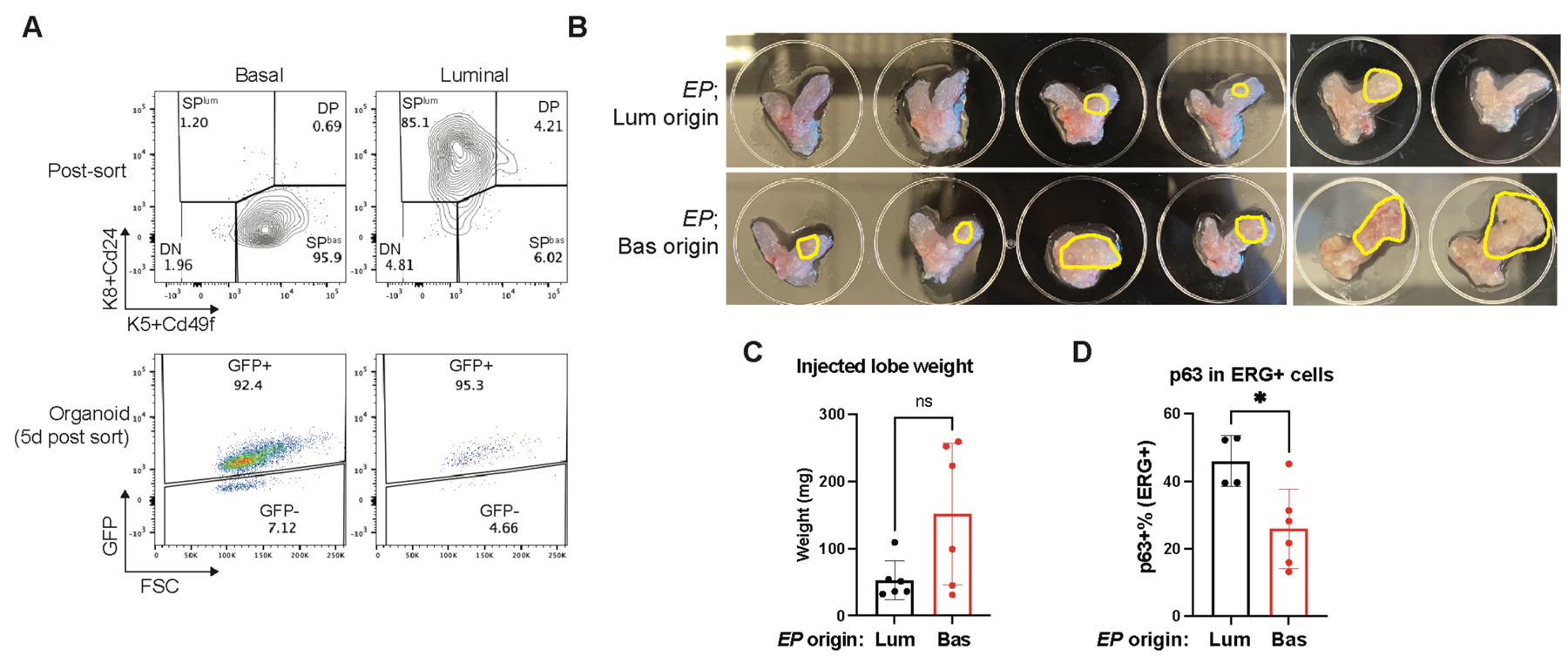
Generation and characterization of *EP* orthografts using freshly isolated and recombined cells. (**A**) Quality control for freshly isolated basal and luminal-derived cells generated in **Fig. 3A**. (Top) Post-sort analysis to validate purity of the sorted basal and luminal populations. (Bottom) Assessment of Cre recombination efficiency using freshly derived organoids harvested at 5 days post Cre. GFP was used as a surrogate for ERG expression. (**B**) Images showing prostates harvested at 5 months post transplantation. Visible grafts, highlighted in yellow circles, suggest a higher graft burden from basal-derived *EP* orthografts. (**C**) Basal-derived *EP* orthografts trended towards a higher injected lobe weight than the luminal derivatives at the 5 months endpoint, further corroborating the higher graft burden as shown in **Fig. 3B**. (**D**) Basal-derived *EP* orthografts displayed a reduction of basal marker p63, further corroborating the notion of luminal fate transition as shown in **Fig. 3D**. p63 expression was measured by flow cytometry in ERG+ graft cells harvested at the 5 months endpoint. Data represent mean ± s.d.; n = 6 (except in **A** where n = 1); ns, not significant; *p<0.05; unpaired two-tailed t-test.

**Figure S7.**
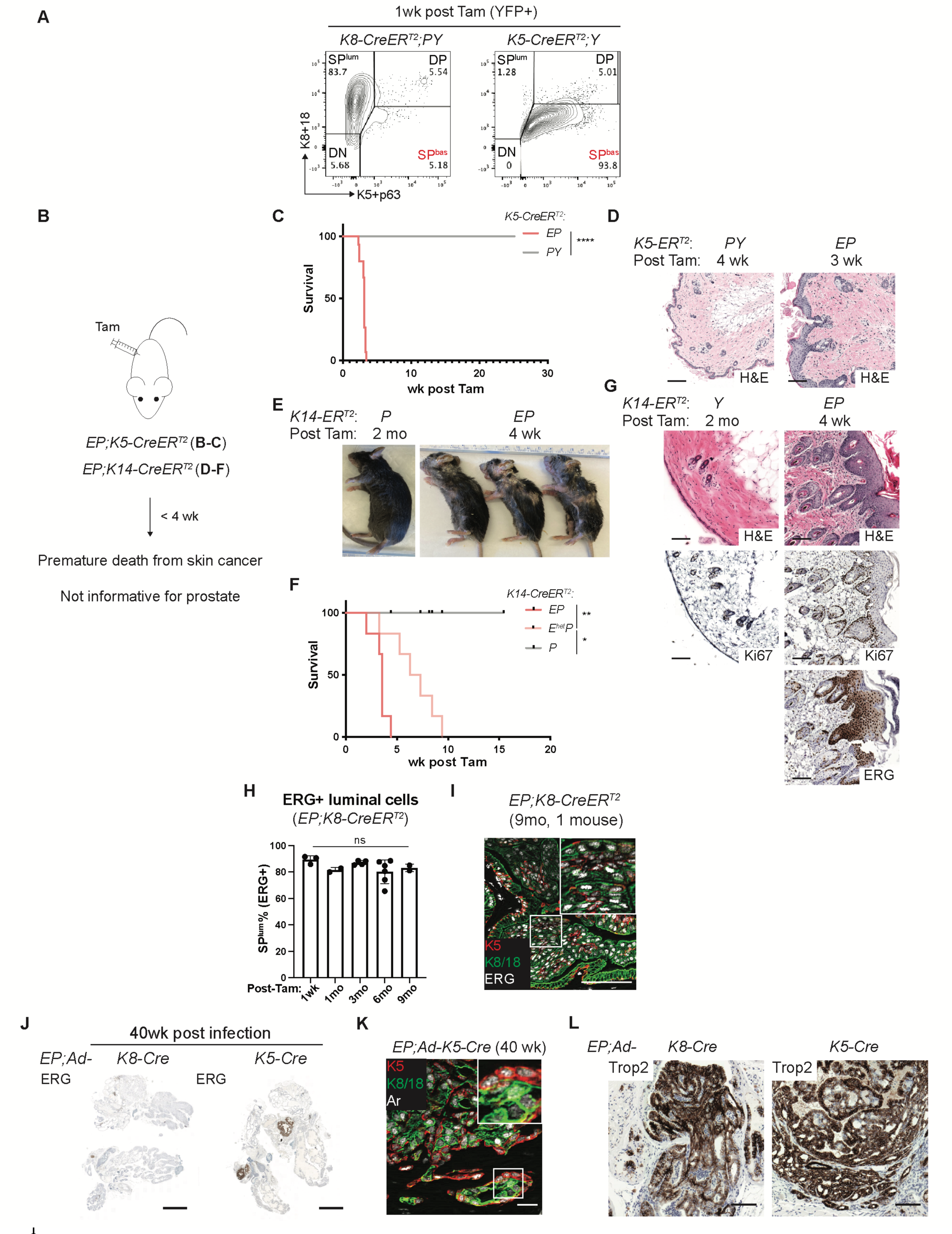
Lineage tracing comparing EP activation from basal and luminal cells. (**A**) Characterization of the cell type specificity of *K5-* and *K8-CreER^T2^*. The basal (SP^bas^) and luminal (SP^lum^) cells identity of the YFP+ recombined cells was measured by flow cytometry at 1 week post tamoxifen administration. (**B**) Schematic summarizing that lineage tracing using *K5* and *K14-CreER^T2^* to activate *EP* leads to a severe skin disease and premature death. This precluded scoring any phenotype in the prostate. (**C-D**) Survival curve (**C**) and skin histology using *K5-CreER^T2^* highlighting an ERG-dependent skin cancer which led to early mouse death. Scale bar, 100µm. (**E-G**) Mice at harvest (**E**), survival curve (**F**) and skin histology (**G**) using *K14-CreER^T2^* highlighting an ERG-dependent skin cancer which led to early mouse death, similar to the *K5-CreER^T2^* model in **C-D**. The severity of the phenotype also correlated with ERG dosage. Ki67 IHC is shown to access the cell proliferation activity. Scale bar, 100µm. (**H**) Flow cytometry showing that ERG+ cells maintain luminal identity in *EP;K8-CreER^T2^*mice through 9 months of tracing. (**I**) IF image showing rare foci of K5+ IM cells from one mouse at the 9 months endpoint. These cells were not clearly identified as a distinct population from flow cytometry in **Fig. 3F**, consistent with the rarity of these cells. Inset shows high-power view. Scale bar, 100µm. (**J**) Sagittal view of whole prostates with ERG IHC highlighting a marked increase in tumor volume in *EP* mice upon Ad-K5-Cre treatment relative to Ad-K8-Cre. Two injection sites are shown. Scale bar, 2mm. (**K**) IF showing Ar expression in both K5- and K5+ cells from invasive adenocarcinomas of *EP;Ad-K5-Cre* mice; inset shows high-power view. Scale bars, 20 µm. (**L**) Trop2 IHC showing a strong Trop2 positivity of the ERG+ lesions, suggesting that the tumor luminal cells acquire a L2 identity^35,36,138^. Trop2 was only detected in the invagination tips in the neighboring glands from adjacent sections from the same mice as in **Fig. 3K**. Note that Trop2 expression was also seen in *EP;Ad-K8-Cre* injected mice, indicative of *EP* activation, but in cells that are not capable of progressing to invasive adenocarcinoma. Scale bar, 100µm.

**Figure S8.**
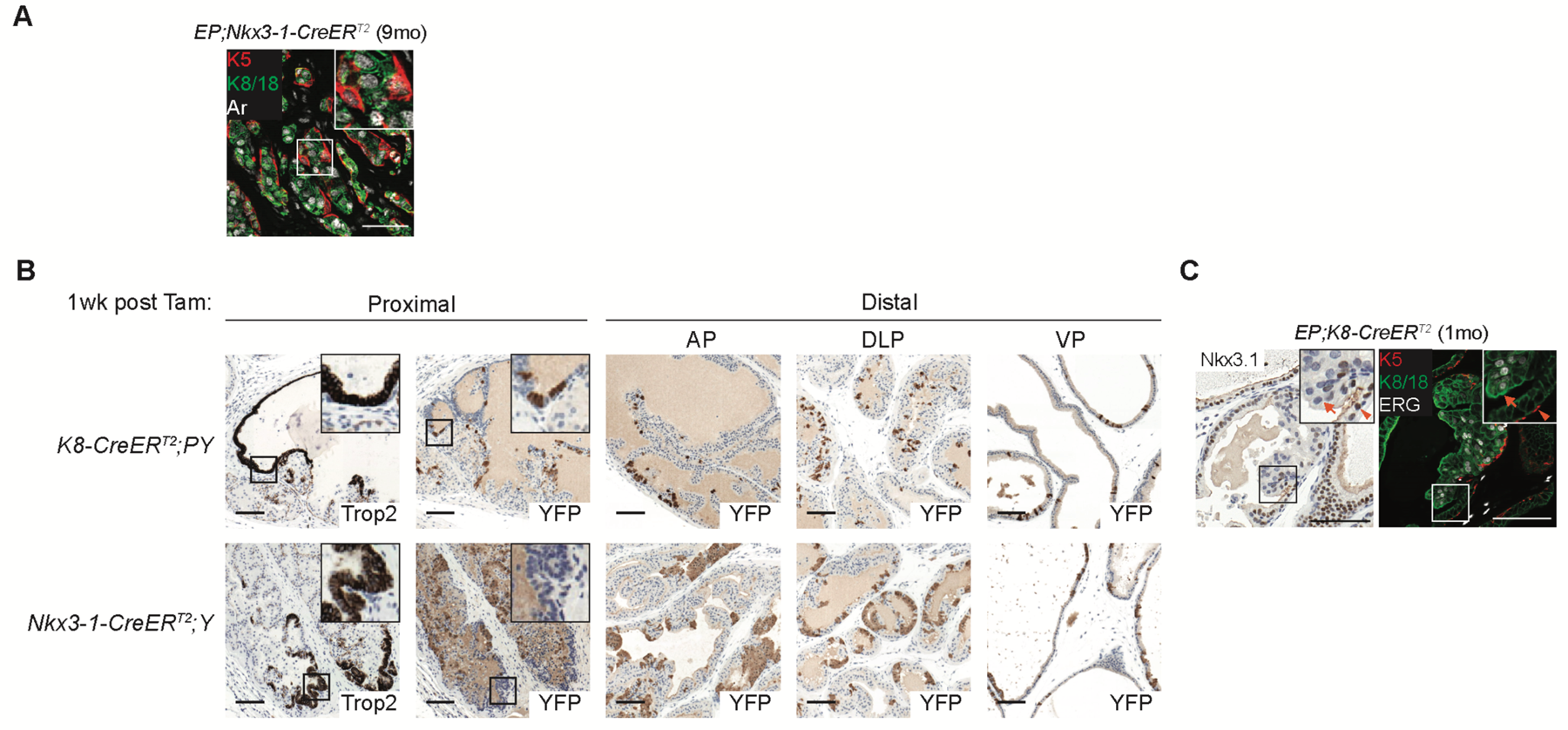
In situ analysis of lineage marker expression and Cre recombination efficiency. (**A**) IF staining showing Ar expression in both K5- and K5+ cells from invasive adenocarcinomas of indicated mice; inset shows high-power view. Scale bars, 50 µm. (**B**) IHC documenting recombined cells after crossing with *K8-CreER^T2^* vs *Nkx3-1-CreER^T2^*mice, YFP was used as a surrogate for Cre recombination. Scale bars, 100 µm. (**C**) *EP;K8-CreER^T2^* mice displayed Nkx3.1 loss in ERG+ cells from the precursor lesions by 1 month post tamoxifen. High-power view in insets highlights the absence of Nkx3.1 signal in ERG+ cells (arrow) and the neighboring Nkx3.1+ normal luminal cells. The *Nkx3-1-CreER^T2^* allele causes haploinsufficiency of the tumor suppressor gene *Nkx3-1^139^* which could potentially explain the tumorigenic phenotypes in *EP;Nkx3-1-CreER^T2^* mice in **Fig. 4**. However, the data here show an early loss of Nkx3.1 in *EP;K8-CreER^T2^*mice as well (potentially reflecting an L2 transition and thus loss of the L1 marker Nkx3.1). Thus, *Nkx3-1* expression is similarly disrupted in both settings and therefore unlikely to cause the different phenotypes. Scale bars, 100 µm.

**Figure S9.**
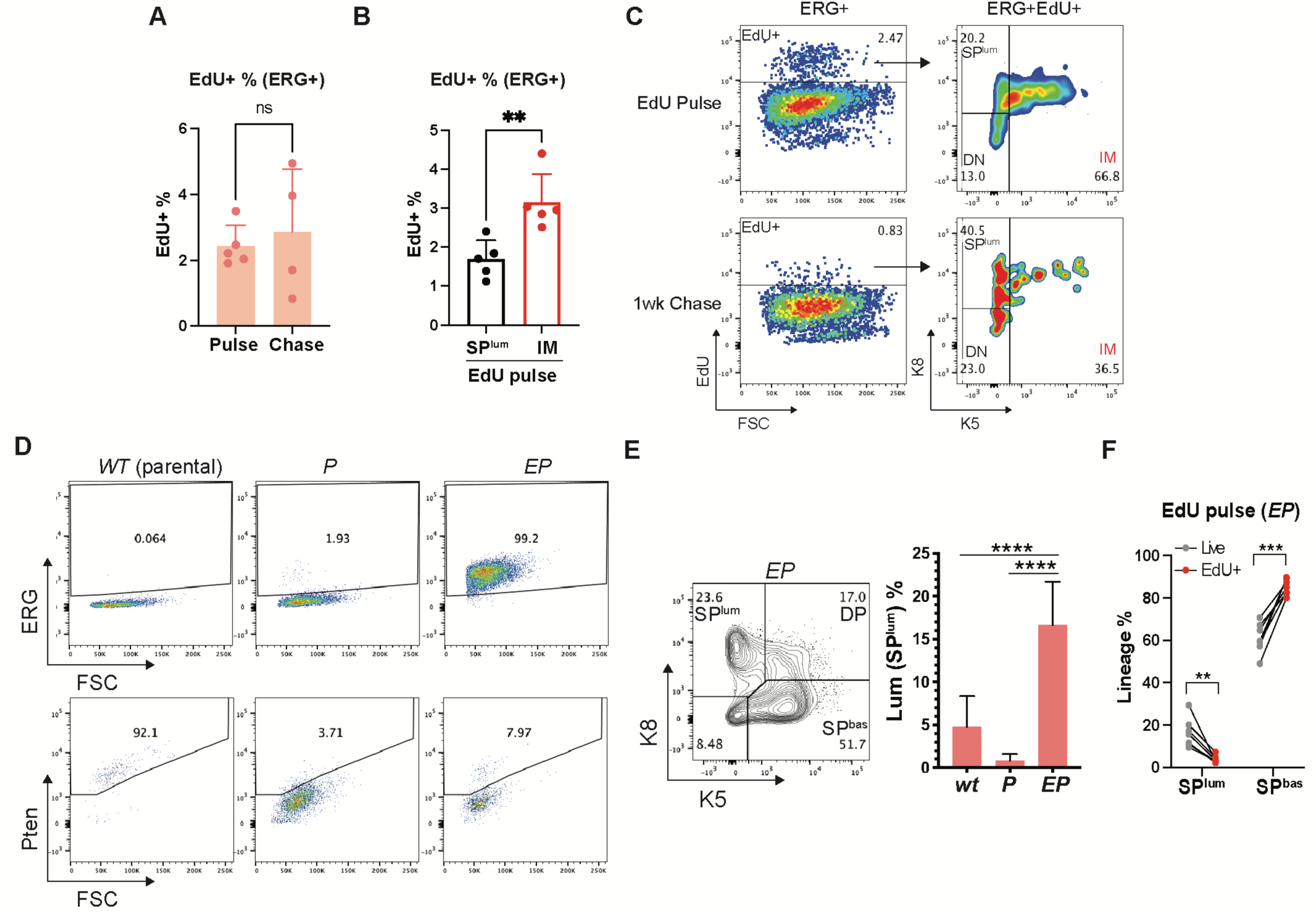
ERG+ basal and intermediate cells proliferate towards a luminal fate *in vitro* and *in vivo*. (**A**) *In vivo* EdU pulse chase assay showing EdU quantification in ERG+ *EPC* cells. The percent of EdU-labeled cells were comparable between the pulse and chase samples, suggesting that the EdU+ population is not likely to enrich the label-retaining cells within 1 week of chase. (**B**) The same samples from **A** showing EdU quantification in SP^lum^ and IM populations of ERG+ *EPC* cells in pulse samples. IM cells showed a higher EdU signal, further corroborating the more proliferative feature of these cells as revealed in **Fig. 1**. (**C**) Flow cytometry analysis on ERG+ cells from **A** highlighting a SP^lum^ shift in EdU+ cells after 1 week of chase. (**D**) Flow cytometry validating the expected ERG and Pten expression status in *EP* organoids and the isogenic controls. (**E**) Flow cytometry showing an ERG-dependent expansion of luminal cells (SP^lum^) in *EP* organoids. (**F**) Flow cytometry quantification in *EP* cells after a EdU pulse. The EdU+ population showed a depletion of luminal (SP^lum^) cells and an enrichment of basal (SP^bas^) cells relative to the total live population. Data represent mean ± s.d.; n > 3.; ns, not significant, *p < 0.05; **p < 0.01; ***p<0.001; ****p < 0.0001; unpaired two-tailed t-test (**A, B, E**); multiple paired t-test with FDR correction by Benjamini, Krieger and Yekutieli (**F**).

**Figure S10.**
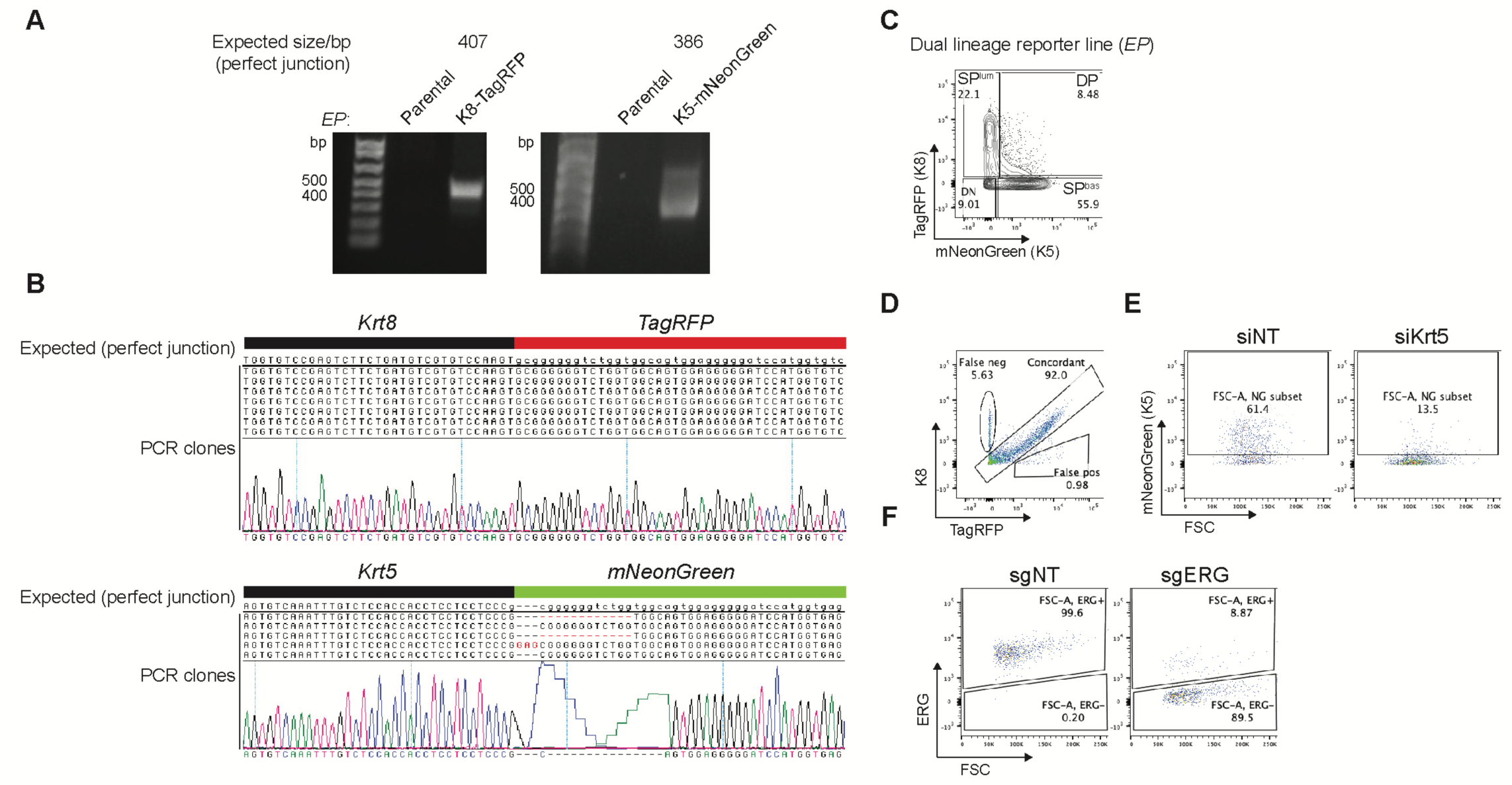
Generation and characterization of the dual lineage reporter knock-in system in *EP* organoids. (**A**) Junction PCR across the target loci and the engineered reporters in the bulk organoid population. The PCR products specific to the knock-in allele with the expected size indicate successful reporter targeting. (**B**) Sanger sequencing of the junction PCR clones from **A** validates expected junction sequences with homogeneous perfect junctions for *Krt8-TagRFP* targeting, and heterogenous junctions with small in-frame insertion/deletions for *Krt5-mNeonGreen* targeting. (**C**) Live cell flow cytometry using the engineered reporter signals recapitulated a similar basal/luminal lineage pattern created by K5/K8 intracellular flow in **Fig. S9E**. (**D**) Intracellular flow cytometry comparing the expression pattern between the endogenous K8 and the TagRFP reporter signal in targeted organoid population. ∼92% of overall concordance was observed. ∼5.6% false-negative population was also observed which either reflects a lack of targeting or a disruption of protein function in a minority of cells. A similar assay was not possible with the *Krt5-mNeonGreen* targeting due to the epitope ablation by the targeting event. (D) Functional validation of the *Krt5-mNeonGreen* engineering by Krt5 depletion, which led to a reduction of the mNeonGreen reporter signal. (**F**) Effective ERG ablation by CRISPR in *EP* reporter organoids. ERG was detected by intracellular flow cytometry 2 days after introducing the indicated CRISPR-RNP.

**Figure S11.**
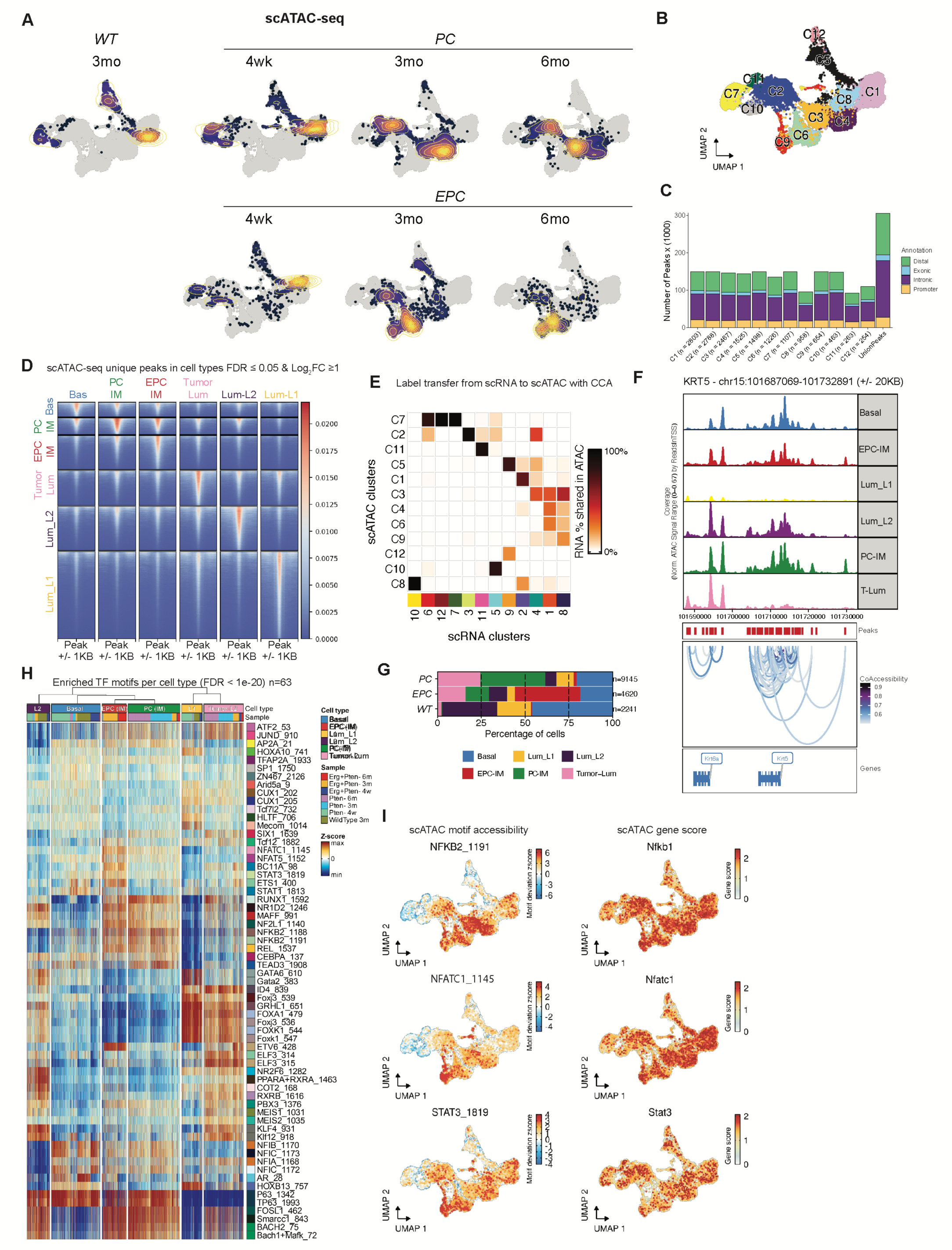
ERG drives a unique chromatin state in intermediate cells. **(A)** scATAC-seq UMAPS by indicated genotypes and time points. Cell density is overlaid and colored by increasing density, from dark to light. **(B)** Clustering generated by leiden algorithm is shown labeled and also colored on scATAC-seq UMAP. **(C)** Barplot showing number of peaks annotated as distal, exonic, intronic, or promoter by cluster. **(D)** Tornado plot showing significantly enriched peaks (FDR <= 0.05 and Log2FC >= 1) for each annotated cell type (Bas, PC-IM, EPC-IM, Tumor-Lum, Lum_L2, Lum_L1) vs all other cell types. Signal plotted is normalized scATAC-seq aggregated by cell type, centered at peak and extended by 1000 nt up and down. **(E)** Grid showing amount of RNA shared with scATAC-seq by cluster. Labels from scRNA were transferred to scATAC using Seurat’s CCA. **(F)** scATAC-seq reads piled up per cell type around the mm10 *KRT5* locus, a classical basal cell marker (top panel). As expected, signal in Lum_L1 is low compared to Basal. Second panel from top shows tick marks to indicate detected peaks. Third panel from top show inferred Co-Accessibility using Cicero, an algorithm used to predict DNA interactions from scATAC-seq data. Fourth panel from top shows *Krt5* gene locus and neighboring *Krt6a* locus in mouse. **(G)** Barplot percentage of cell type per genotype is shown. **(H)** scATAC-seq heatmap of most enriched transcription factor candidates when comparing one cell type to all others. Cells are grouped by cell type as columns and rows indicate motifs from database. The heatmap elements are colored by chromvar inferred z-score. Motif significance was calculated (see methods) and those having an p-value < 1e-20 after BH-false discovery rate correction are shown. **(I)** Representative UMAPs colored by motif deviation z-score (left) and inferred gene expression (gene score, right) of EPC-IM enriched TFs (NFKB2, NFATC1, STAT3).

